# A BODIPY-naphtholimine-BF_2_ dyad for precision photodynamic therapy, targeting and dual imaging of endoplasmic reticulum and lipid droplets in cancer

**DOI:** 10.1101/2024.01.07.574513

**Authors:** Nitish Chauhan, Mrunesh Koli, Rajib Ghosh, Ananda Guha Majumdar, Ayan Ghosh, Tapan Kumar Ghanty, Soumyaditya Mula, Birija Sankar Patro

## Abstract

Currently effective therapeutic modalities for pancreatic ductal adenocarcinoma (PDAC) is not available, leading to gloomy prognosis and ~6-months median patient survival. Recent advances showed the promise of photodynamic therapy (PDT) for PDAC patients. Next generation photosensitizers (PS) are based on “organelle-targeted-PDT” and provides new paradigm in the field of precision medicines to address the current challenge for treating PDAC. In this investigation, we have constructed a novel PS, named as **N*b*B**, for precise targeting of endoplasmic reticulum (ER) and lipid droplets (LD) in PDAC, as malignant PDAC cells are heavily relying on ER for hormone synthesis. Our live cell imaging and fluorescence recovery after photobleaching (FRAP) experiments revealed that **N*b*B** is instantly targeted to ER and LD and show simultaneous dual fluorescence colour due to polar and non-polar milieu of ER and LD. Interestingly, the same molecule generates triplet state and singlet oxygen efficiently and cause robust ER stress and apoptosis in two different PDAC cells in the presence of light. Together, we present, for the first time, a potential next generation precision medicine for ER-LD organelle specific imaging and PDT of pancreatic cancer.

## INTRODUCTION

Pancreatic ductal adenocarcinoma (PDAC) is one of the most difficult to treat cancers with dismal and grim prognosis rate, due to unavailability of therapeutic modality.

^[1,2]^ Pancreatic cells are heavily involved in hormone, digestive enzyme synthesis and secretion, with an extensive network of endoplasmic reticulum (ER).^[3]^ Besides, malignant cells rely on higher protein synthesis requirements and ER functionality than normal cells.^[4]^ ER is a hub for different signaling pathways and acts as a double-edged sword at the interface of cell survival and death. ER is also involved in the generation of lipid droplets (LD), which in turn positively correlated with advanced clinical staging, metastasis, and poor survival. ^[5,6]^ These facts make the ER and LD as attractive targets for sensitizing malignant PDAC cells.^[7]^

Although multiple approaches failed, a growing number of studies however have indicated that photodynamic therapy (PDT) may be a viable approach for treatment of pancreatic tumors. Especially clinical trials with PDT agents, mesotetrahydroxyphenylchlorin and verteporfin, showed some positive therapeutic outcomes for PDAC patients.^[8]^ However, considering the fact that next generation photosensitizers (PS) are target specific, ER-organelle-targeted-PDT of PDAC is not yet explored. Development of such precision photomedicines may address the unmet challenge of PDAC treatment.

For targeting and imaging of ER and LD, few interesting recent reports showed (i) imaging of ER^[9-11]^ or LD^[12-16]^ with multiple probes and (ii) simultaneous imaging of ER and LD with one single probe but with one emission colour, ^[17-18]^ leading to inherent difficulties in discriminating structure and functions of ER and LD. To the best of our knowledge, only one elegant study on a single probe (3-hydroxyflavone system) based simultaneous imaging of ER and LD with dual emission colour was reported ^[19]^. 3-hydroxyflavone system, however, lack photosensitization properties for selective killing of cancer cells. In this regard, we report for the first time a single agent with multiple functionalities: (1) high ER and LD specific targeting, (2) imaging ER and LD with dual fluorescence and (3) robust photosensitizing ability (Figure 1A). We and others have shown that BODIPY-based systems are promising design choices individually for cellular imaging, photosensitizations and targeting.^[20-22]^ For rationale designing and functional tuning, we have chosen “Naphthalene Scaffold” for following attributes. (i) **ER targeting**: Naphthalene based various natural and synthetic drugs target ER efficiently.^[23-25]^ (ii) **Simultaneous ER-LD dual imaging**: Since, ER and LD have major water and lipid contents, respectively in their core, we anticipated that naphthalene moiety, reported to have excellent self-aggregation properties ^[26,27]^, may influence the fluorescence emission of BODIPY backbone in ER and LD. (iii) **Photosensitizing properties:** One of the key requirements for the PDT is generation of singlet oxygen (^1^O_2_) through energy transfer from excited triplet state (T_1_) of PS to molecular oxygen. In general, BODIPYs are highly fluorescent with very low triplet quantum yields. Although BODIPY substituted with iodine/bromine are reported for efficient ^1^O_2_ generation for PDT ^[28,29]^, toxic heavy metals are least preferred in drugs or biological studies. Orthogonal BODIPY dimers are also efficient triplet PSs. A recent preliminary study has shown meso-substituted BODIPY with naphthalene make them good PS, which generate T_1_ and ^1^O_2_ efficiently.^[30]^ Moreover, meso- and *β*-substitutions have a profound effect on the excited state dynamics of the BODIPY. ^[31,32]^ However, nothing is attempted yet to explore the photophysical, photochemical and photobiological roles of dyads, where modified napathalene is conjugated to BODIPY in the form of naphtholimine boron complex (naphtholimine-BF_2_).

**Figure 1.**
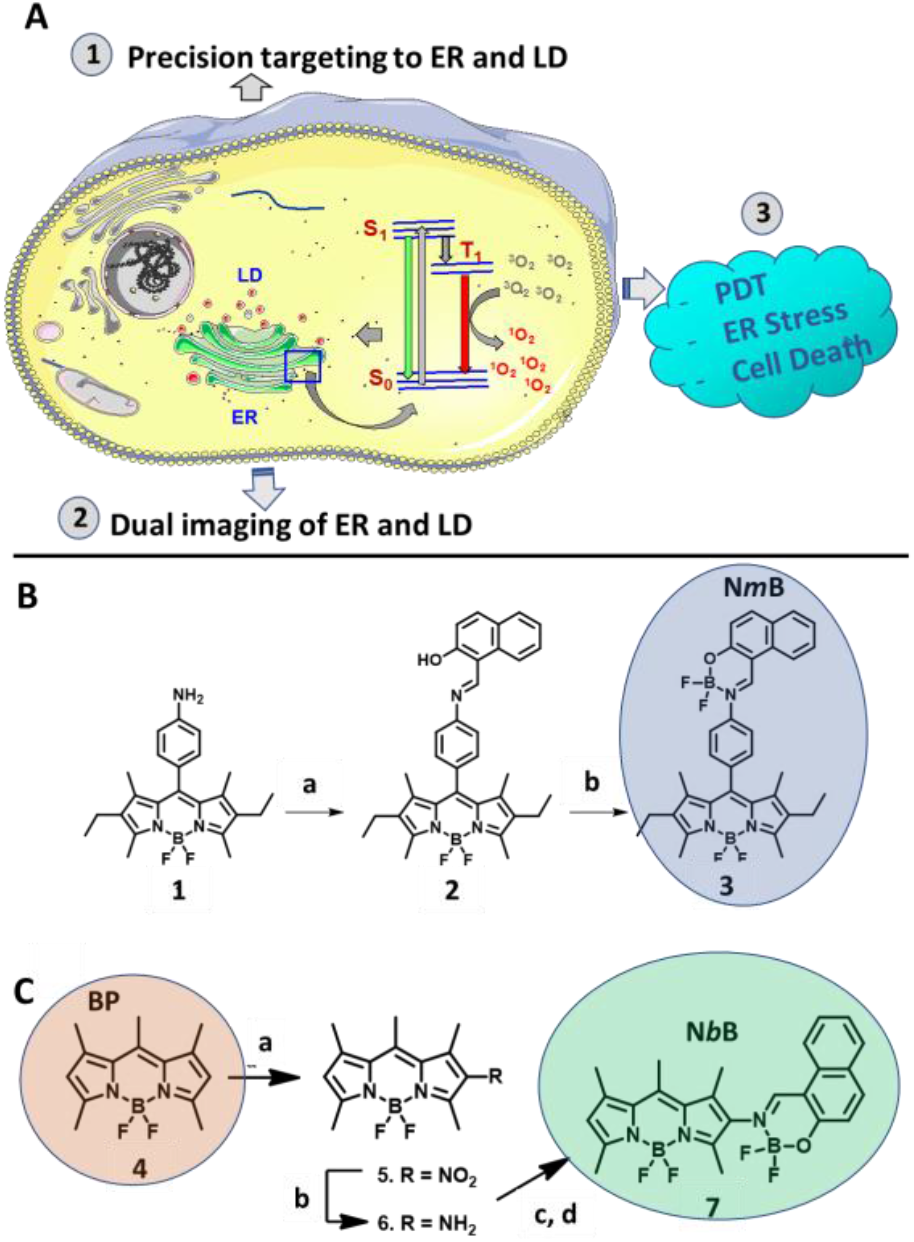
Targeting and synthesis of molecules for ER specific PDT. (A) Potential of a single molecule for targeting, dual imaging of ER and LD and photosensitization of cancer. (B) Synthesis routes of **N*m*B**: a) 2-Hydroxy-1-naphthaldehyde, dry MeOH, reflux, 4 h; b) Et_3_N, BF_3_·OEt_2_, 25 °C, 24 h. (C) Synthesis routes of **N*b*B**: a) HNO_3_, 0 °C, 30 min; b) H_2_, 10% Pd/C (10 %), CH_2_Cl_2_/EtOH (1:1), 25 °C, 24 h; c) 2-Hydroxy-1-naphthaldehyde, PTS (cat. Amount), Na_2_SO_4_, DCE, 70 °C, 12 h; d) Et_3_N, BF_3_.OEt_2_, DCE, 70 °C, 12 h.

## RESULTS AND DISCUSSION

Taking clues from the above reports, we rationally designed and synthesized a new class of *bis*-chromophoric fluorescent dyad by fusing naphtholimine-BF_2_ at *<u>m</u>eso*-linked <u>B</u>ODIPY (**N*m*B**) and *<u>β</u>*-linked <u>B</u>ODIPY (**N*b*B**) (Figure 1B, C). For synthesis, dye **1** was condensed with 2-hydroxy-1-naphthaldehyde to generate Schiff base **2**, which was treated with excess BF_3_·OEt_2_ to furnish **3 (N*m*B)**. Further, dye **4** (**BP**) was nitrated (**5**) and hydrogenated to obtain 2-amino BODIPY (**6**). Dye **6** was condensed with 2-hydroxy-1-naphthaldehyde to form corresponding Schiff base, which was treated with excess BF_3_·OEt_2_ to furnish **7 (N*b*B)**. The detailed synthesis and characterization of the reaction products are shown in Supplementary information (Figure S1-S6).

UV/Vis absorption, fluorescence spectra and photophysical properties of the **NmB, N*b*B** and **BP** were evaluated in dicholoromethane (DCM) (Figure 2A, S7, Table S1). Narrow absorption bands, high molar extinction coefficients (ε_max_), high fluorescence quantum yield (Φ_fl_) with small Stokes shifts (*ν*) were the spectral characteristics of all the three dyes. Interestingly, both **N*m*B** and **N*b*B** dyes showed greenish yellow fluorescence but with drastically compromised intensity (Φ_fl_ **=** 0.04 for **N*m*B** and Φ_fl_ **=** 0.09 for **N*b*B**), as compared to **BP** (Table S1). It is important to mention here that the Stokes shift of the dye **N*b*B** is ~3 times higher than the **BP** (*Vide infra* for explanation). Strikingly, the **N*b*B** shows negative solvatochromism, *i*.*e*., hypsochromic shift of the fluorescence maxima with minimal effects on absorption maxima in different solvents with increasing dielectric constant.^[33,34]^. The λ_max (em)_ of **N*b*B** was 514.2 nm in water, which is red-shifted by ~29 nm in cyclohexane (λ_max (em)_ = 543.6 nm). In contrast, this shift was only ~6 nm for **N*m*B** dye (Figure 2B, S8, Table S2, S3), suggesting that linking of naphtholimine-BF_2_ derivative at *β*-position of BODIPY may be critical and have profound effect on the self-aggregation-based shift of fluorescence emission of **N*b*B** dye in non-polar environment. This conclusion was further supported by the observation that yellow fluorescence of **N*b*B** in DCM solution turns red due to self-aggregation in solid state (Figure 2C, D).

**Figure 2.**
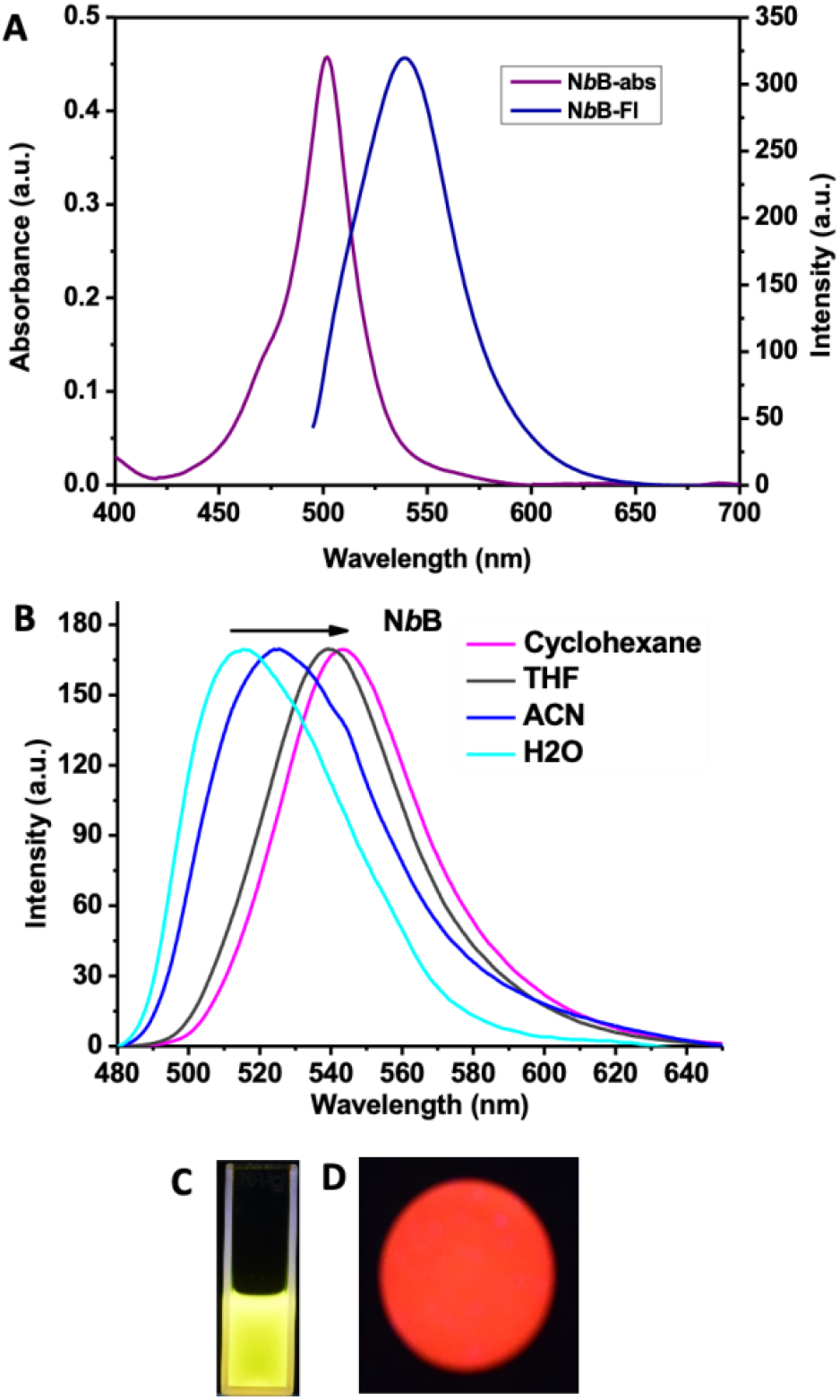
Photophysical properties of **N*b*B**. (A) Absorbance and fluorescence spectra of **N*b*B** in DCM. (B) Fluorescence spectra of **N*b*B** were acquired in different solvents. (C) Fluorescence of **N*b*B** in solid state (KBr pallet).

Considering a significant effect of solvent polarity on λ_em_, it is expected that naphthaolimine moiety in **N*b*B** dye may direct it for (1) ER-specific localization and (2) allow change of intrinsic fluorescence, when it is channelized from ER (polar core) to LD (non-polar core). In order to evaluate such phenomenon, MIA-PaCa-2, pancreatic ductal adenocarcinoma cells were incubated with **BP, N*m*B** and **N*b*B** (30 min) and visualized under confocal laser scanning microscope. The green fluorescence (λ_ex_ = 488 nm) of all three dyes was exclusively localized to the cytoplasm (Figure 3A) and the intensity was in the order of **BP**>**N*b*B>N*m*B** (Figure 3B). Although, fluorescence emission of **N*b*B** was lower as compared to **BP**, it is sufficiently high for microscopic studies. Interestingly, **N*b*B** can simultaneously label two separate sub-cellular structures with dual colours, as some portion of “green fluorescence” also displayed intense “red fluorescence” in the form of punctae (Figure 3A, B). In contrast, **N*m*B** and **BP** displayed no such properties (Figure 3A). Further, cellular green fluorescence of **N*b*B** showed excellent co-localization with commercially available ER-specific probe (ER-Tracker Blue) while its co-localization was poor with mitochondria (Mito-Tracker Red) and lysosome specific probe (Lyso-Tracker Red) (Figure S9), confirming an efficient ER-targeting effect when naphtholmine-BF_2_ derivative is conjugated at *β*-position of **BP**. Besides, **N*b*B** stained red fluorescing punctae also colocalized precisely with lipid droplets, globular organelles with high refractive index, in differential interference contrast image (DIC) (Figure 3B).^[19]^ We further employed Lambda Scanning Mode (LSM) in confocal microscopy, a powerful tool employs single λ_ex_ for characterizing fluorescence emission spectrum of the dye in different cellular organelles/locations. LSM acquisition at λ_ex(488nm)_ revealed that fluorescence emission of **N*b*B** was evident in aqueous-(polar)-core of ER in the range of 505-565 nm (λ_max_ = 531.8 nM; Green region) and significantly drops at >570 nm (Figure 3D, E). Interestingly, fluorescence emission of **N*b*B** was retained significantly in nonpolar core of LD in 570-638 nm region (δ_max_ between ER and LD = 585 nm; Yellow region; Figure 3D, E). These fluorescence spectra of **N*b*B** in ER (aqueous) and LD (nonpolar) region are in excellent agreement with its photophysical properties in solution (Figure 2B).

**Figure 3.**
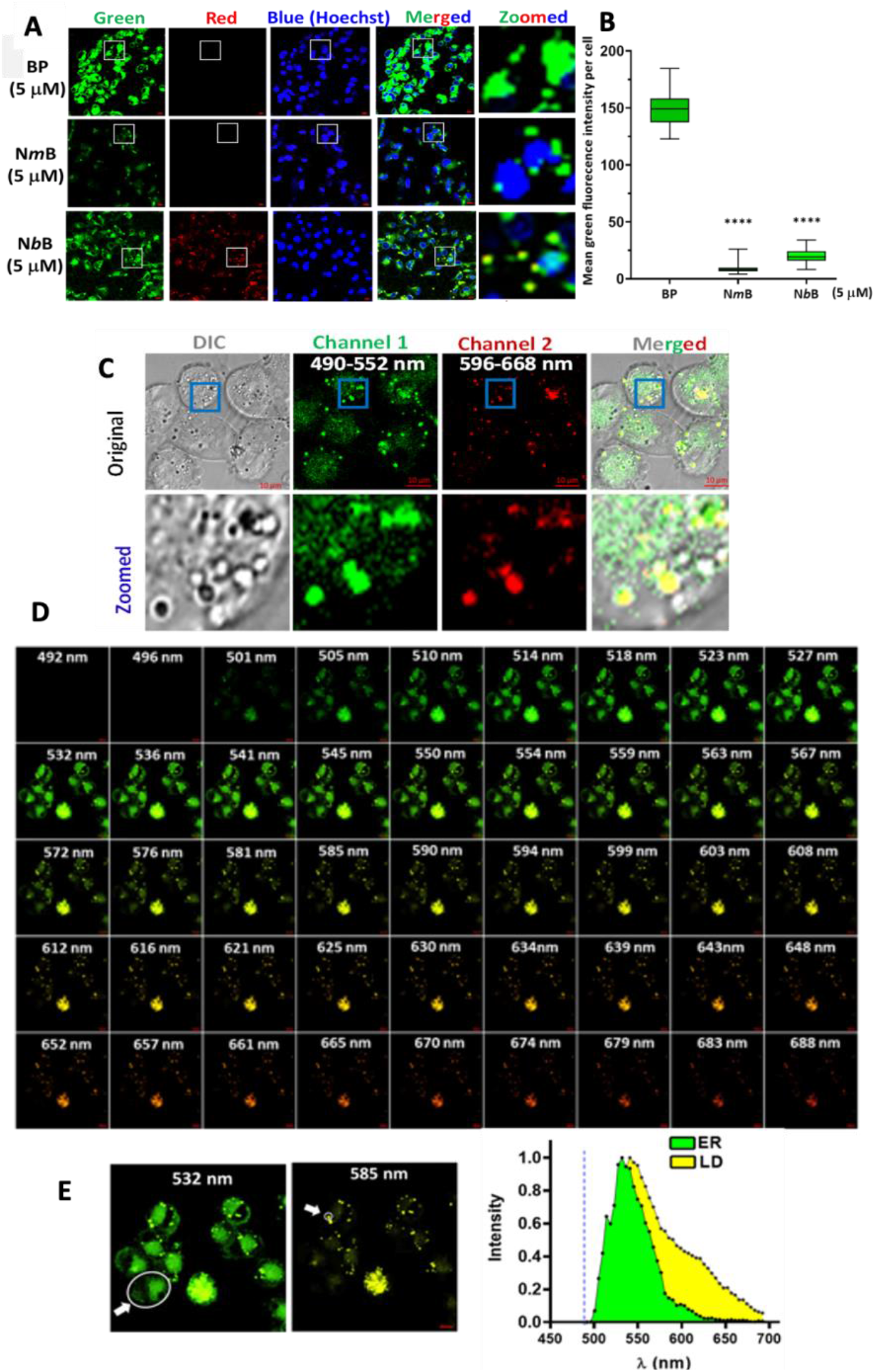
Localization and dual fluorescence properties of **N*b*B** in ER and LD in cancer cells. (A, B) MIA PaCa-2 cells were treated with **BP, N*m*B** and **N*b*B** (5 µM each) for 30 min and fluorescence images were acquired in confocal microscope. Scale bar = 20 µm. Mean green fluorescence intensity per cell in channel 1 (green) was quantified in different dye staining. \*\*\*\**p<0*.*0001* w.r.t **BP** treatment. (C) MIA PaCa-2 cells were treated with **N*b*B** (5 µM) for 30 min and DIC and fluorescence images were acquired in multi-channel mode confocal microscope. Zoomed images show localization of **N*b*B** in ER (green only) and LD (yellow colour due to colocalization of green in channel 1, red in channel 2 and black in DIC channel for globular lipid droplets). (Channel 1: λ_ex_ = 488 nm; Channel 2: λ_ex_ = 594 nm). (D, E) *In situ* fluorescence emission of **N*b*B** (5 µM) in MIA-PaCA-2 cells, as acquired in λ-scan mode. Enlarged images showing ER and LD specific dual emissions of **NbB** at 532 and 585 nm are shown in “E”. Arrows indicate the ROI markings for ER and LD, respectively. Fluorescence spectra at the ROI at ER and LD (Dashed line in the graph = λ_ex_ at 488 nm).

To further confirm that *in situ* dual florescence properties of **N*b*B** is dependent on the differential polarity of solvent in ER and LD, we have performed confocal microscopy based FRAP (fluorescent recovery after photobleaching) assay.^[35]^ In this assay, **N*b*B** fluorescence in the small region of interest (ROI) in ER and LD were photobleached through micro-irradiation with high-intensity laser. Subsequently, uptake/diffusion of the **N*b*B** from cytoplasm (polar milieu) to ER and LD was assessed measuring the kinetics of fluorescence recovery in respective ROIs in live cell imaging. We observed a quick and complete recovery of **N*b*B** fluorescence (green) in ROI in ER (Figure 4A, B). Contrarily and intriguingly, we observed that (1) the fluorescence recovery (yellow fluorescence) was severely affected and (2) appeared in green in colour in LD (ROI), suggesting the diffusion of cytoplasmic aqueous **N*b*B** solution (green) into the LD (ROI) without being mixed with nonpolar core of the LD (Figure 4A-C). Together, these results showed a first case of design of a novel synthetic molecule for dual targeting and imaging of cellular ER and LD. Of note, these properties of **N*b*B** are not specific to one PDAC cell line, as similar ER-LD targeting were also obtained with another PDAC cells e.g., PANC-1 (Figure S10).

**Figure 4.**
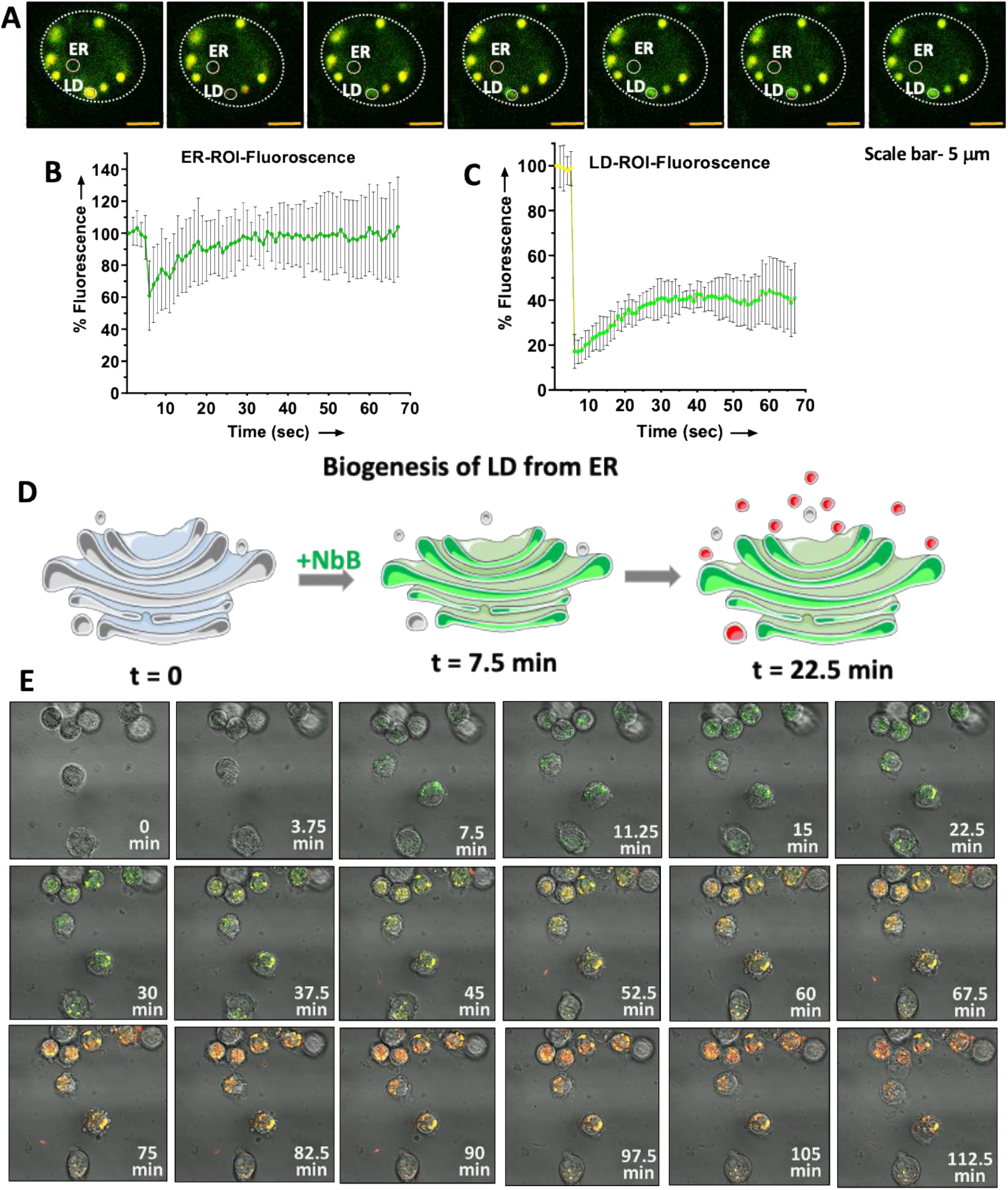
Dual fluorescence of **N*b*B** is associated with different polarity in ER and LD (A-C) Florescence recovery after photobleaching (λ_ex_ = 488 nm) in ROI in ER and LD in MIA PaCa-2 cells were measured, after labelling of cells with **N*b*B** (5 µM, 30 min). (D, E) Scheme and images showing the kinetics of labelling of **N*b*B** (5 µM) in ER and LD organelles in MIA PaCa-2 cells.

On further characterization, we found that **N*b*B** could able to dual label both ER and LD distinctly at a concentration as low as 0.5 µM (Figures S11, S12). In order to understand how **N*b*B** is routed to two different cellular organelles (ER and LD) and their possible interdependence, we followed the kinetics of cellular localization of **N*b*B** in MIA-PaCa2 cells, employing live cell confocal imaging. Our data revealed that the fluorescence of **N*b*B** promptly appeared at ER (green) within 4 min of staining, subsequently yellow-red coloured LD emerged from the ER (Movie M1, Figure 3D, E). Moreover, formation of yellow-red LD particles was almost confined to ER (green) region, supporting the previous report that biogenesis of LD particles is originated from ER through retrograde ER-LD trafficking process.^[19]^

After establishing first two properties (ER targeting and dual imaging), we ventured to explore the PDT effects of these novel conjugates. One of the critical requirements of the PDT is efficient T_1_ formation and subsequent ^1^O_2_ generation for killing cancer cells. The ^1^O_2_ generation capacities was determined by monitoring the dye-sensitized photooxidation of 1,3-diphenylisobenzofuran (DPBF) (see supplementary method). Under visible light photoirradiation (>495 nm), ***NbB*** induced ^1^O_2_ mediated photooxidation of DPBF, which was assessed by the rapid decay of absorption peak (412 nm) of DPBF (Figure 5A). Imperatively, the order of ^1^O_2_ generation/photooxidation capacity was **N*b*B**>**N*m*B**>**BP** (Figure S13), suggesting that the triplet conversion of the **N*b*B** is very high *vis-à-vis* **N*m*B** and **BP**. Further, to assess the extent of T_1_ formation of all three dyes, nanosecond flash photolysis was used (see methods). As shown in Figure 5B, a strong triplet-triplet absorption signal (425 nm) and very strong bleach at 500 nM was observed for **N*b*B**, while weak/no signal was observed for **N*m*B** and **BP** (7 ns pulse, Ex_532 nm_). Moreover, this signal of **N*b*B** decays faster in air (life time: 0.65 μs) than in nitrogen atmosphere (life time: 37 μs) (Figure 5C, D), suggesting efficient transfer of energy from T_1_ to generate ^1^O_2_. The triplet yield of the **N*b*B** was estimated to be ~90 (±5)% by comparing the bleach signal with platinum octaethylporphyrin, a standard triplet sensitizer, under identical experimental conditions (Figure S14). Despite low fluorescent quantum yield of **N*m*B** and **N*b*B**, only later has the ability to generate high levels of T_1_ and ^1^O_2_. This anomalous behavior was further explained by theoretical calculations. By using B3LYP/def-TZVP level of theory, we observed that electron density being completely shifted towards *meso*-naphtholimine-BF_2_ moiety in LUMO (lowest occupied molecular orbital) or BODIPY core in HOMO (highest occupied molecular orbital), due to torsion angle in **N*m*B** (Figure 5E). Besides, electron withdrawing nature of *meso*-naphtholimine-BF_2_ alters the localization of HOMO and LUMO and hence enhances the nonradiative emission coupled weak fluorescence of **N*m*B**. ^[36]^ In contrast, we did not see any charge redistribution as both HOMO and LUMO are located mainly on the BODIPY moiety and slightly spread over the *β*-naphtholimine-BF_2_ moiety in S_0_ ground state geometry of **N*b*B** (Figure 5F). Further, its *β*-naphtholimine-BF_2_ and the BODIPY segment are not coplanar and bear a dihedral angle (C11-N12-C16-C17 = 73.6^o)^ between them (Figure 5F). However, the *β*-naphtholimine-BF_2_ moiety in the S_1_ excited state was almost orthogonal with the BODIPY core (C11-N12-C16-C17 = 96.7°) (Figure 5G). This geometry relaxation on photoexcitation increases the Stokes shift considerably.^[36]^ This explains and supports the fluorescence quenching of **N*b*B** is due to its conversion to T_1_ state. Together, above results confirms the unique ability of **N*b*B** dye for high T_1_ and ^1^O_2_ yield, which makes **N*b*B** also an excellent PDT candidate.

**Figure 5.**
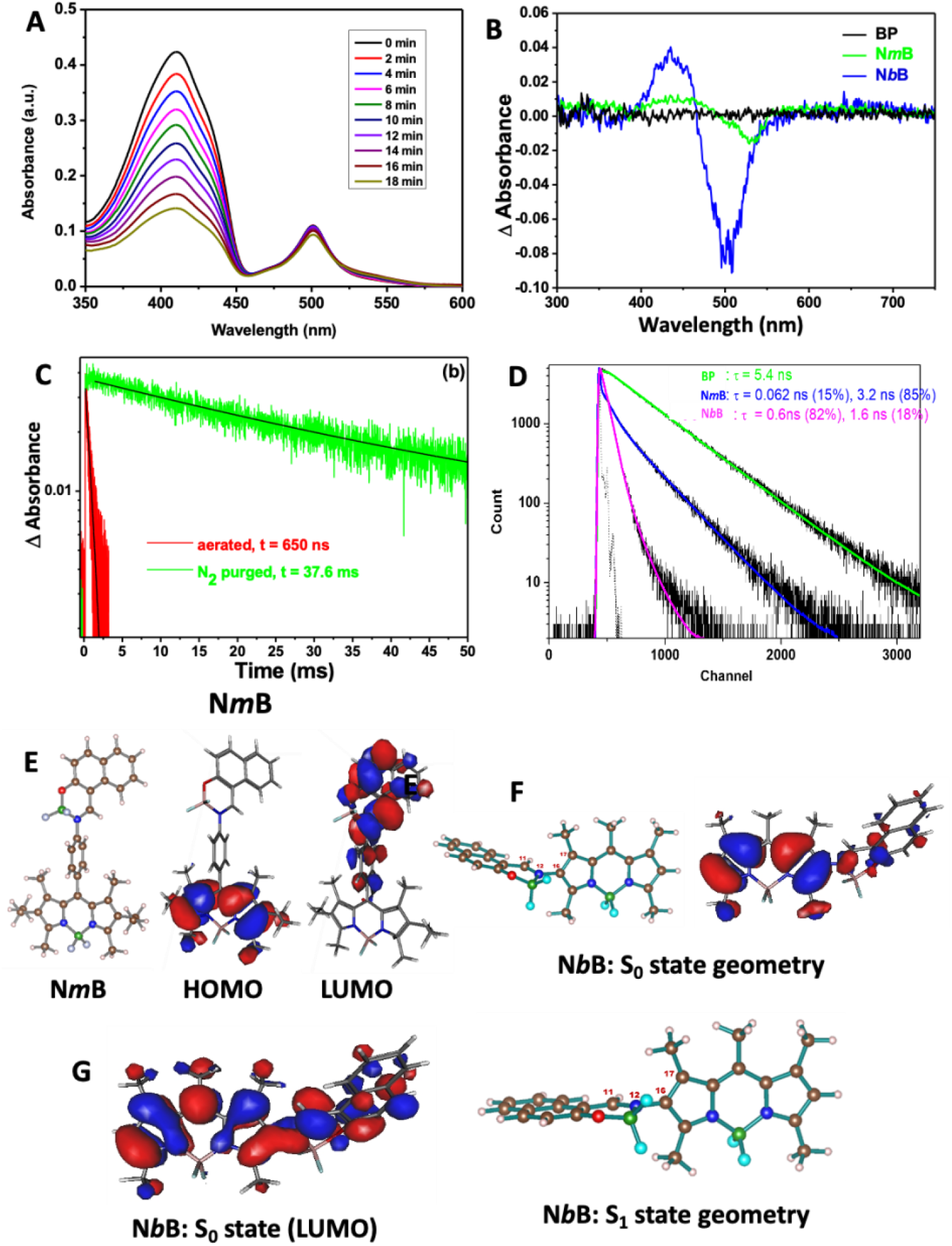
(A) **N*b*B** (5 µM) sensitized photooxidation of DPBF (50 µM) was measured in term of change in absorption spectrum of DPBF during photoirradiation (light source: >495 nm). (B) Nanosecond transient absorption spectrum of dyes **BP, N*m*B** and **N*b*B** and (C) triplet decay kinetics of **N*b*B** in dichloromethane (λ_ex_ = 532 nm, λprobe = 425 nm). (D) Emission decay traces (at 550 nm) of the dyes **BP, N*m*B** and **N*b*B** in dichloromethane were measured by TCSPC method. Solid lines are the exponential fit to the experimental data and obtained lifetimes are given in the inset. (E, F) Optimized structure of **N*m*B** and **N*b*B**.

Considering the above impressive properties of **N*b*B** (ER-LD targeting and high T_1_ and ^1^O_2_ generation), we sought to know whether **N*b*B** can also able to kill PDAC by selectively targeting ER-LD through PDT. As shown in Figure 6A, our results revealed that both **BP** and **N*m*B** were almost completely failed to reduce colonogenic growth of MIA PaCa-2 cells in the absence/presence of light. Interestingly, **N*b*B** could reduce clonogenic growth in a concentration dependent manner (dark toxicity at 7.5 µM), which was robustly photo-induced in two different PDAC cells (MIA PaCa-2 and PANC-1; Figures 6B, C, S15A, B). At lower concentrations, dark toxicity of the **N*b*B** was significantly supressed while PDT mediated killing was still apparent in two different PDAC cells (MIA PaCa-2 and PANC-1). A strong synergistic interaction between **N*b*B** and light was observed for PDT mediated killing of both PDAC cells, as assessed by Comebenefit software (Figures 6D, S15D). Similar results were also observed for short term MTT assay (Figure S16A, B). These results are in agreement with the photochemical properties of the **N*m*B, N*b*B** and **BP** (Figure S13). Mechanistically, the PDT effect of **N*b*B** is mediated through induction of sub-G1 and cleavage of caspase-3, the markers of apoptosis, in PDAC cells (Figure 7A-D, S17). Since **N*b*B** targets ER and has the ability to generate copious amount of ^1^O_2_ in the presence of light, PDT effects of **N*b*B** might be attributable to ER stress. Recent reports show that ER stress culminates to ROS (reactive oxygen species) generation and glutathione (GSH) depletion, leading to cell death.^[37,38]^ We could not able to assess the generation of cellular ROS in **N*b*B** mediated PDT process, as fluorescence spectrum of the **N*b*B** interferes with commercially available ROS analysing dyes (DCFDA, DHE etc). Alternatively, we have analysed GSH level by monobromobimane (MBB), which emits blue florescence after reacting with cellular GSH. In this regard, treatment of MIA PaCa-2 and PANC-1 cells with **N*b*B** led to significant reduction of cellular GSH in a time dependent manner (Figure 7E, F, S18A-C, S19A-C). In contrast, **N*b*B** or light alone treatment failed to induce such significant changes in PDAC cells. Imperatively, PDT treatment with **N*b*B** significantly affected the expression of ER-stress markers e.g. IRE-α and BiP levels were increased while Calnexin level was reduced (Figure 7G, H). These observations confirmed “targeted-PDT effect” of **N*b*B** dye on ER of the PDAC cancer cells.

**Figure 6.**
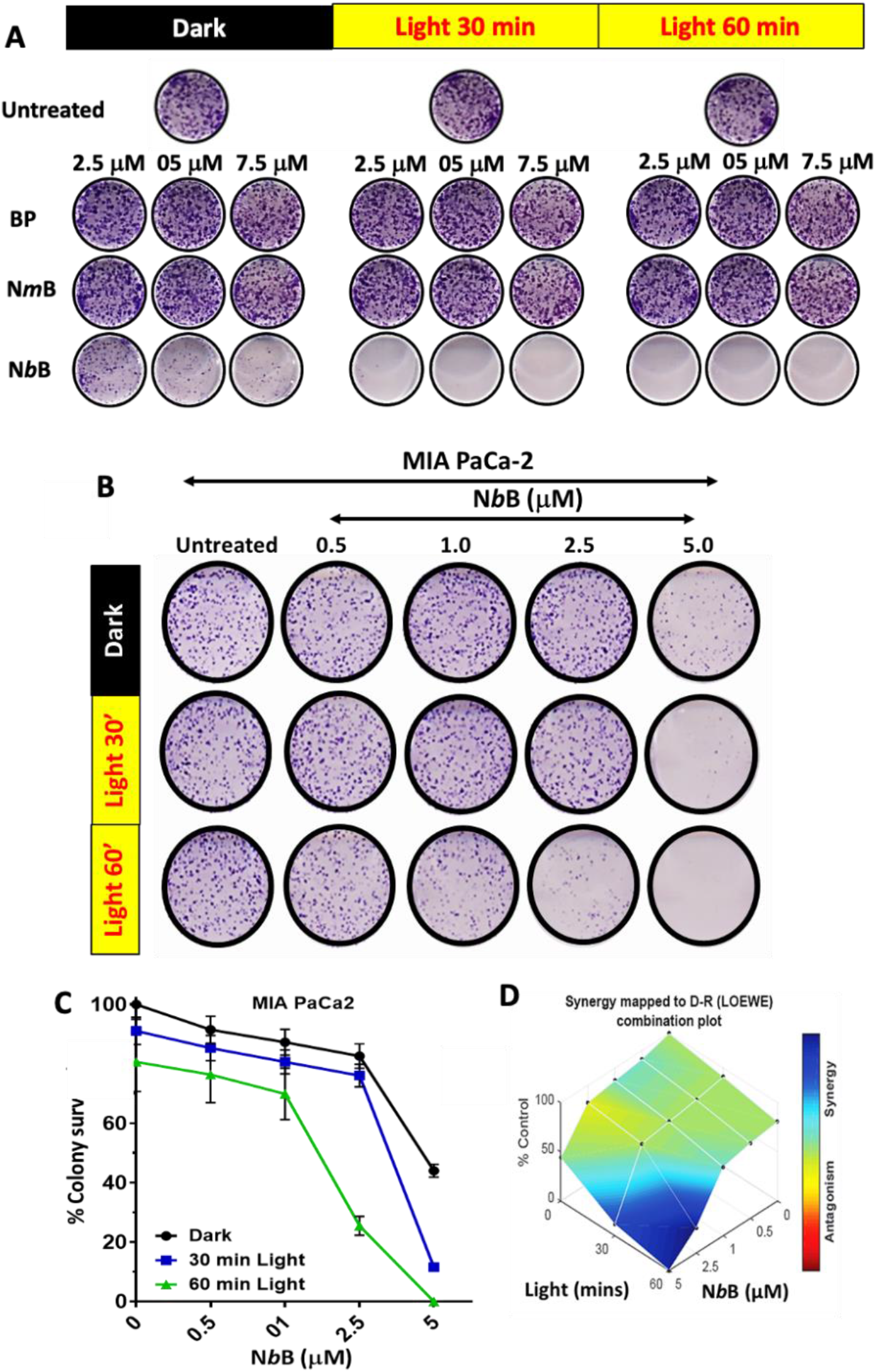
**N*b*B** efficiently photosensitises pancreatic cancers. (A) MIA PaCa-2 cells were treated with different concentration of **BP, N*m*B** and **N*b*B** for 30 min and exposed to visible light. Clonogenic survival was assessed after 9-10 days of treatments. (B-D) MIA PaCa-2 cells were treated with **N*b*B**, as mentioned above, at lower concentrations and clonogenic survival was assessed. Representative images of clonogenic assay are shown in “B”. Quantification for the colony growth and Comebenfit based synergism (**N*m*B** and light) are shown in “C” and “D”, respectively. (C, D) NbB induces apoptosis (sub-G1, Caspase-3 activation, 24 h) during PDT effects on MIA-PaCa-2 cells. (E) PDT effects of NbB is mediated through robust activation of ER stress markers.

**Figure 7.**
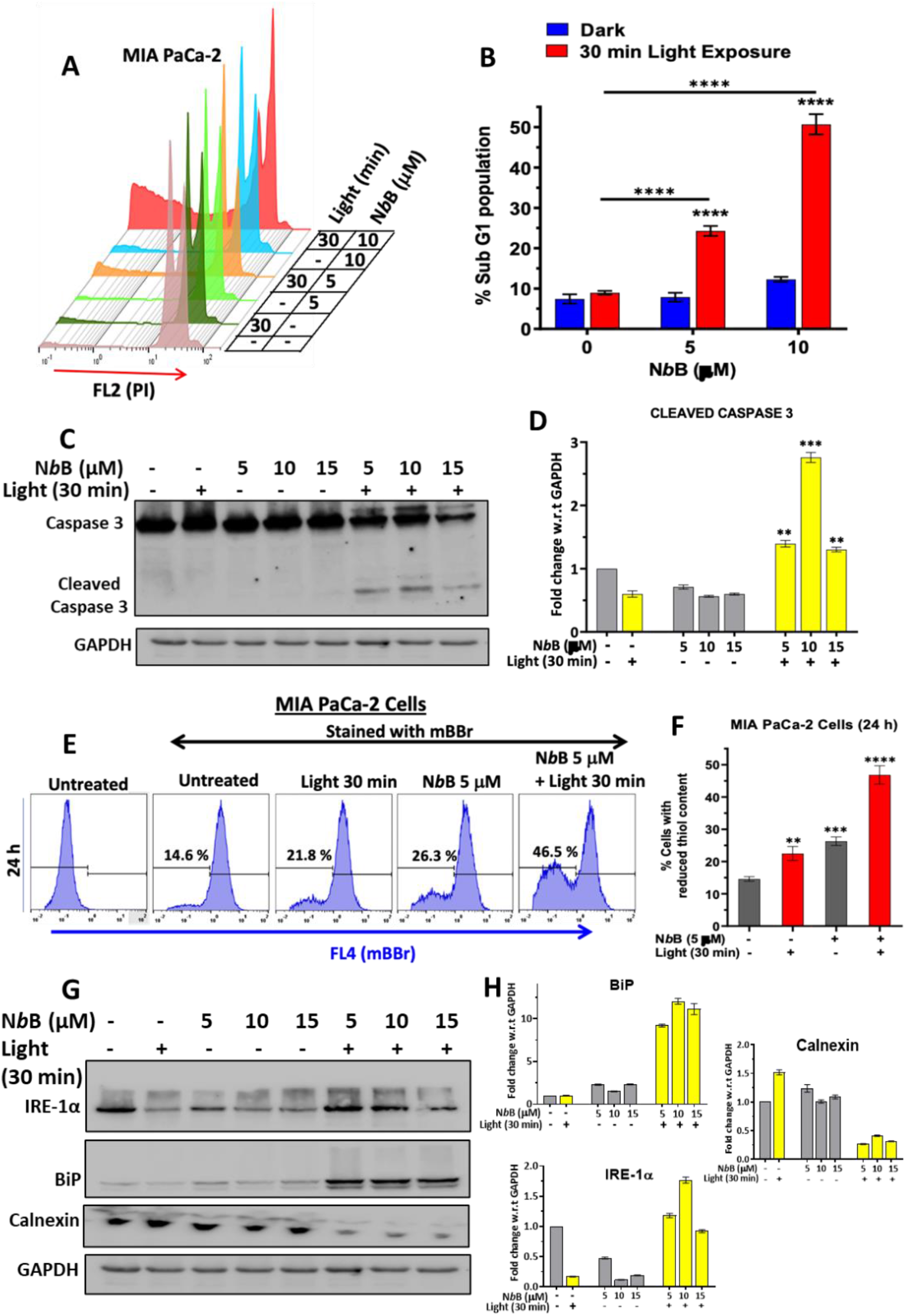
**N*b*B** induces ER stress and apoptosis in pancreatic cancers. (A) MIA PaCa-2 cells were treated with different concentration of **N*b*B** for 30 min and exposed to visible light. Apoptosis was assessed by sub-G1 assay after 24 h of treatments. Quantifications for % sub-G1 in different treatments are shown in “B”. (C, D) MIA PaCa-2 cell were treated with **N*b*B** as above and activation of caspase-3 was assessed and quantified. (E, F) MIA PaCa-2 cell were treated with **N*b*B** as above and cellular thiol level was assessed by MBB based flow cytometry assay. (G, H) PDT effects of **N*b*B** is associated through robust activation of ER stress markers. MIA PaCa-2 cell was treated with **N*b*B** as above and ER stress markers are assessed by Western blotting.

## CONCLUSIONS

In conclusion, this work illustrates for the first time a novel BODIPY based dyad with three biological functions: (1) targeting ER and LD, (2) dual imaging of ER and LD and (3) potent PDT effects for sensitization of pancreatic cancer. Till date an efficient photosensitization of PDAC through precise targeting of ER and LD in cancer is not known. Based on several reports, PDT is emerging as a viable approach for treatment of pancreatic tumors.^[39]^ Moreover, advanced fiber optic based light delivery has been facilitated the PDT treatment at pancreatic sites.^[39]^ This work also underscores the importance of a novel *bis-* chormophoric molecule - (naphtholimine-BF_2_ and BODIPY conjugate) for targeting ER in PDAC, aggregation induced fluorescence emission shift and robust and efficient triplet state and ^1^O_2_ generation. Since, PDAC cells are over-reliant on ER network for hormonal secretion, comprehensive targeted PDT effect on ER may help in developing efficient therapy against pancreatic tumors.

## MATERIALS AND METHODS

### Materials

For organic synthesis, all the chemical reagents and solvents were purchased from a local supplier and were used without any further purification. Fetal bovine serum (FBS), Antibiotic-Antimycotic solution (#15240096), Dulbecco’s Modified and Eagle Medium (DMEM) were procured from Gibco, 3-(4,5-dimethylthiazol-2-yl)-2,5-diphenyltetrazolium bromide (MTT, #M5655), propidium iodide (PI), crystal violet, RNase, Hoechst 33342, monobromobimane (MBB) and anti-GAPDH (#G8795) were from Sigma-Aldrich Chemicals. MitoTracker Red, ER-Tracker Red, and Lyso-Tracker Red are from Invitrogen. Anti-caspase-3 (#CST-9662), anti-IRE-1α (#CST-3294), anti-BiP (#CST-3177) and Calnexin-C5C9 (#2679) were purchased from Cell Signalling Technology. Lumi-Light ECL-plus western blotting kit was from Roche Applied Science.

### Synthesis and Characterization

A detail protocol for the synthesis of the studied molecules is mentioned on the supporting information.

### Photophysical Properties

Measurement of fluorescence lifetime of the samples dissolved in desired solvents were carried out by time correlated single photon counting (TCSPC) technique using IBH-UK (Horiba-Jobin-Yvon) system. 511 nm laser light with a FWHM of about 100 ps was used for excitation of samples and the emission kinetic traces were collected at emission peak. The kinetic traces were fitted exponentially using the IBH DAS 6.2 software module. Triplet spectra and kinetics of the samples were measured by laser flash photolysis technique using a laser kinetic spectrometer (Edinburgh Instruments, U.K.; model LP920). Samples were excited by a 7 ns Nd:YAG laser at 532 nm (second harmonic of 1064 fundamental laser) and probed with a 450 W pulsed xenon lamp. All spectroscopic measurements were carried out at room temperature.

### Cell Culture

Pancreatic Ductal Adenocarcinoma (PDAC) cell lines MIA PaCa-2 and PANC-1, purchased from the European Collection of Authenticated Cell Cultures (ECACC), were maintained as per the instruction provided by ECACC. The cell lines were cultured in DMEM having 2 mM glutamine, supplemented with 10% Fetal bovine serum (FBS) and 1% Antibiotic-Antimycotic solution. Cells were grown in a 5% CO2 and 95% humidity environment at 37 °C temperature. All the experiments were performed within 6-8 passages from the hawing of a cell line from liquid nitrogen.

### Photo-exposure Protocol

The cells were irradiated with white light from a 100W incandescent bulb source. The distance between the surface of irradiated cells and the bulb was 25 cm. The photoirradiation was done for two different time points (30 and/or 60 min).

### Clonogenic Assay

Clonogenic assay was performed for the evaluation of the cytotoxic effect of the compound, following report^[40]^ with minor modifications. Briefly, cells were seeded (0.8 x 10^3^/well) in 6 well plates in a complete medium for overnight. Cells were treated with **N*b*B** and light as mentioned above and incubated for 8-10 days. Post incubation, cells were washed with PBS, colonies were fixed with chilled methanol for 10 minutes, and colonies were stained with a 0.5 % (w/v) crystal violet solution (30 min). Staining solutions were discarded and washed 3-4 times with water. Stained colonies containing more than 30 cells each were counted under an inverted light microscope.

### Confocal Microscopy

Cells were seeded on a glass coverslip (1 x 10^5^ cells/coverslip) for overnight. Cells were incubated with different concentrations of florescent dyes (**N*m*B, N*b*B** and **BP**) for the indicated time periods. Post-incubation cells were washed twice with PBS and then mounted with 80 % glycerol before imaging through the confocal microscope (LSM780, Carl Zeiss). For **N*m*B, N*b*B** and **BP**, images were acquired by laser excitation at 488/543/594 nm while Hoechst 33342 was excited with 355 nm laser. For colocalization studies, cells were incubated with **N*b*B** (05 µM) and LysoTracker Red (100 nM) or MitoTracker Red (200 nM) or ER-Tracker Red (500 nM) or ER-Tracker Blue (300 nM) for 20 min. Post-incubation cells were washed twice with PBS and then mounted with 80 % glycerol before imaging through the confocal microscope (Zeiss, LSM780). ER-Tracker Red, MitoTracker Red, LysoTracker Red or ER-Tracker Blue was excited using 594/355 nm laser. Fluorescence intensity and colocalization parameters were assessed using Zen software (Carl Zeiss).

### Live Cell Imaging and In Situ Lambda Scan Imaging

Live cell imaging and lambda mode scanning of the stained cells were carried out with the confocal microscope (LSM 780, Carl Zeiss), which has an in-built cell incubation chamber (5% CO2 and 95% humidity) for carrying out live cell imaging experiments for long hours. Cells were seeded on to the thin glass bottom (35 mm) of confocal dishes (SPL Life Sciences #100350) for overnight. Cells were treated with NbB (5 µM) and kinetics of appearance of fluorescence of NbB, from t0 to t115-min at the interval of 3.75 min, in cells were acquired in channel 1 (λ_ex_ = 488 nm; λ_em_ = 490-552 nm) and channel 2 (λ_ex_ = 594 nm; λ_em_ = 596-668 nm). For in situ lambda scan imaging, cells were treated as above in confocal dishes and spectral imaging in lambda scan mode was carried out in LSM 780 microscope. The NbB dye was excited at 488 nm and spectral imaging (fluorescence emission) was acquired at every 4 nm from 492-688 nm.

### Fluorescence Recovery After Photobleaching (FRAP) Assay

This assay was carried out as per our previous report for FRAP experiments ^[35]^, with minor modifications. Briefly, Cells were seeded on to the thin glass bottom of confocal dishes for overnight. Cells were treated with **N*b*B** (5 µM, 30 min) and FRAP assay was carried out in LSM 780 confocal microscope. Images were captured using a 6x digital zoom and a 40x magnification to make sure a 1 ms pixel dwell time. The region of interest (bleach ROI in ER and LD) was bleached using 100% power of 488 nm laser line for 40 repetitions after five pre-bleach photos (488 nm laser line, 1% power) were taken. In addition, photobleaching was taken into consideration during image capture by using a reference region of interest. A total of 100 pictures were taken after bleaching at intervals of one millisecond. Utilizing Zen Blue 3.4 software, the % recovery of **N*b*B** fluorescence was measured after picture acquisition. The formula shown below was employed to determine the percentage of fluorescence recovery:

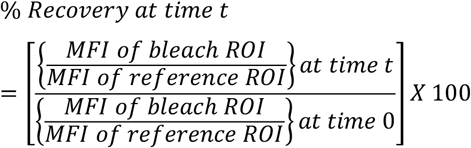

MFI stands for mean fluorescence intensity in this context. Data on percentage recovery was used to generate recovery graphs.

### Dye Sensitized Photo-oxidation Study ^[39]^, Synergistic interaction Study (Comebenefit) ^[41]^, MTT ^[42]^, sub-G1 ^[43]^, Cellular Protein Thiol Estimation ^[44]^ and Western Blotting ^[41]^

These assays are performed as per the previous reports. Detailed protocols with minor modifications are included in the Supporting Information.

### Statistical Analysis

At least three independent experiments were carried out, and each individual experiment was done in triplicate. GraphPad Prism 8 was used to create all the generated graphs and conduct the statistical analysis. The statistical significance of the data was examined using the unpaired t-test and analysis of variance (ANOVA). Significant was defined as a probability value of p<0.05. Cells that had been exposed to the vehicle were regarded as untreated controls.

## ASSOCIATED CONTENT

### Supporting Information

The Supporting Information is available free of charge on the Bioarxive website. Detailed experimental procedures and characterization: ^1^H NMR, ^13^C NMR, HRMS; additional figures (PDF)

## AUTHOR INFORMATION

### Author Contributions

^‡^NC and MK contributed equally. NC, MK, SM and BSP conceptualized and designed experiments. MK and SM synthesized the molecules and performed photophysical studies. RG performed nanosecond transient studies. AG and TKG performed the theoretical studies. NC, AGM and BSP executed the biological experiments. BSP prepared and finalized the manuscript after taking inputs from all the authors.

### Notes

The authors declare no competing financial interests.

## ACKNOWLEDGMENT

This work is supported financially by Department of Atomic Energy, India.

