## Supporting Information for "A BODIPY-naphtholimine-BF_2_ dyad for precision photodynamic therapy, targeting and dual imaging of endoplasmic reticulum and lipid droplets in cancer"

### Materials and Methods

**Synthesis and Characterization.** All the chemical reactions were carried under argon atmosphere using anhydrous solvents in screw-cap schlenk tube. Thin-layer chromatography was used to monitor the progress of the reactions using commercially available 0.25 mm fluorescent silica plate (F-254) and visualized using a UV-lamp (254 nm and 365 nm wavelength). Compounds were purified by flash chromatography using silica gel (40-63 µm). NMR spectra were recorded in 800 MHz Bruker FT-NMR & 500 MHz and 600 MHz Varian FT-NMR instruments. High Resolution Mass Spectrometric (HRMS) analysis was done using 6540 UHD Accurate-Mass Agilent Q-TOF LC/MS instrument.

**Synthesis of Schiff base 2 (Scheme 1).** A mixture of **1** (500 mg, 1.26 mmol) and the 2-hydroxy-1-naphthaldehyde (261 mg, 1.51 mmol) in dry MeOH was refluxed for 4 h. The solvent was removed under a reduced pressure and the crude product was recrystallized from $CH_2Cl_2$/MeOH to get corresponding Schiff base **2** as orange red crystals (570 mg, 82%). $^1H$ NMR (600MHz, $(CD_3)_2CO$, 25 °C, TMS; Figure S1): δ 1.00 (t, *J* = 7.2 Hz, 6H), 1.45 (s, 6H), 2.38 (q, *J* = 7.2 Hz, 4H), 2.51 (s, 6H), 7.13 (d, *J* = 9.2 Hz, 1H), 7.39 (t, *J* = 7.4 Hz, 1H), 7.52 (d, *J* = 7.9 Hz, 2H), 7.57 (t, *J* = 7.7 Hz, 1H), 7.81-7.85 (m, 3H), 7. 98 (d, *J* = 9.0 Hz, 1H), 8.51 (d, *J* = 8.5 Hz, 1H), 9.87 (s, 1H); $^{13}C$ NMR (200 MHz, $(CD_3)_2CO$, 25 °C, TMS; Figure S2): δ = 12.2, 12.6, 14.9, 17.5, 110.2, 120.9, 121.0, 121.9, 122.0, 122.5, 124.6, 128.6, 129.0, 130.1, 130.6, 131.6, 133.7, 134.3, 134.6, 137.4, 139.1, 141.1, 147.7, 147.9, 154.5, 158.1, 158.3, 168.6, 169.1 ppm; HRMS (ESI-TOF) m/z: $[M+H]^+$ Calcd. for $C_{34}H_{34}BF_2N_3O$: 550.2836; Found: 550.2841.

**Synthesis of N*m*B (Scheme 1).** A mixture of **2** (100 mg, 0.18 mol) and diisopropylethylamine (0.1 mL, 0.46 mmol) in dry $CH_2Cl_2$ was stirred for 30 min. Then $BF_3{\cdot}Et_2O$ (0.1 mL, 0.45 mmol) was added to the mixture and the solution stirred at 85 °C for 2 h. The resulting dark mixture was washed with aqueous saturated $NaHCO_3$, water, and brine and dried. Removal of solvent in vacuo followed by column chromatography of the residue (silica gel, petroleum ether -EtOAc) furnished **N*m*B**, which was recrystallized from $CH_2Cl_2$/cyclohexane to afford orange needles (100 mg. 92%). $^1H$ NMR (800 MHz, $CDCl_3$, 25 °C, TMS; Figure S3): δ 1.00 (t, *J* = 7.2 Hz, 6H), 1.37 (s, 6H), 2.33 (q, *J* = 7.2 Hz, 4H), 2.55 (s, 6H), 7.33 (d, *J* = 8.8 Hz, 1H), 7.49 (d, *J* = 8.0 Hz, 2H), 7.52 (t, *J* = 8.0 Hz, 1H), 7.69

(t, $J$ = 8.0 Hz, 1H), 7.77 (d, $J$ = 8.0 Hz, 2H), 7.89 (d, $J$ = 8.0 Hz, 1H), 8.14 (d, $J$ = 8.8 Hz, 1H), 8.16 (d, $J$ = 9.6 Hz, 1H), 9.21 (s, 1H); $^{13}$C NMR (200 MHz, $CDCl_3$, 25 °C, TMS; Figure S4): δ = 12.2, 12.7, 14.7, 17.2, 109.0, 119.4, 120.7, 124.4, 125.5, 128.3, 129.9, 130.1, 130.7, 131.6, 133.3, 136.9, 138.3, 141.8, 143.3, 154.5, 158.1, 163.4 ppm; HRMS (ESI-TOF) m/z: $[M+H]^+$ Calcd. for $C_{34}H_{33}B_2F_4N_3O$: 598.2818; Found: 598.2808.

**Synthesis of 4,4-Difluoro-1,3,5,7,8-pentamethyl-2-nitro-4-bora-3a,4a-diaza-s-indecene (Compound 5) (Scheme 1):** Compound **4** (**BP)** (100 mg, 0.38 mmol) was added to cooled (0 °C) $HNO_3$ (8.5 mL, 42%), and the resulting orange mixture was stirred at 0 °C for 1.5 h. The mixture was filtered, washed with $H_2O$ (4 × 10 mL), and dried under vacuum. Column chromatography of the residue (silica gel, hexane/EtOAc) furnished **5** which was recrystallized from $CHCl_3$/hexane to afford orange needles (95 mg, 81.5%) $^1$H NMR (200 MHz, $CDCl_3$, 25 °C, TMS): δ 2.49 (s, 3H), 2.59 (s, 3H), 2.70 (s, 3H), 2.71 (s, 3H), 2.80 (s, 3H), 6.29 ppm (s, 1H); $^{13}$C NMR (50 MHz, $CDCl_3$, 25 °C, TMS): δ 14.1, 14.3, 15.1, 17.6, 18.0, 125.2, 132.0, 136.0, 143.7, 146.8, 147.6, 162.5 ppm; EIMS: m/z (%): 307.1 (26) $[M]^+$, 308.1 (100) $[M+1]^+$; elemental analysis calcd (%) for $C_{14}H_{16}N_3BF_2O_2$ : C 54.75, H 5.25, N 13.68; found: C 54.35, H 5.12, N 13.77.

**Synthesis of 2-Amino-4,4-difluoro-1,3,5,7,8-pentamethyl-4-bora-3a,4a-diaza-s-indecene (Compound 6) (Scheme 1):** $HCO_2NH_4$ (21 mg, 0.33 mmol) and Zn dust (22 mg, 0.33 mmol) were added to a solution of 4,4-difluoro-1,3,5,7,8-pentamethyl-4-bora-3a,4a-diaza-*s*-indecene (86 mg, 0.33 mmol) in MeOH (20 mL), and the mixture was stirred for 10 min until the orange solution turned dark pink. The reaction mixture was eluted through Celite, the eluate was concentrated under vacuum, and the residue was subjected to column chromatography (basic alumina, hexane/ethyl acetate) to furnish **7** (72 mg, 80%) as brown solid. $^1$H NMR (500 MHz, DMSO-$d_6$, 25°C, TMS): δ 2.22 (s, 3H), 2.32 (s, 3H), 2.34 (s, 3H), 2.38 (s, 3H), 2.53 (s, 3H), 6.01 (s, 1H); $^{13}$C NMR (125 MHz, 25 °C, TMS): δ 11.9, 12.4, 13.8, 15.9, 16.4, 117.8, 118.4, 129.9, 131.6, 135.6, 137.3, 139.7, 146.6, 148.8 p pm; EI-MS m/z: 278.2 $[M+1]^+$.

**Synthesis of N*b*B (Scheme 1).** A mixture of **6** (150 mg, 0.54 mmol) and the 2-hydroxy-1-naphthaldehyde (112 mg, 0.65 mmol) in dry MeOH was refluxed for 4 h. The solvent was removed under a reduced pressure and the crude product of corresponding Schiff base was obtained as orange powder which was used for next step. The crude mixtures (200 mg) and diisopropylethylamine (0.18 mL, 1.04 mmol) in dry $CH_2Cl_2$ was stirred for 30 min. Then $BF_3{\cdot}Et_2O$ (0.12 mL, 1.0 mmol) was added to the mixture and the solution stirred at 85 °C overnight. The resulting dark mixture was washed with aqueous saturated $NaHCO_3$, water, and brine and dried. Removal of solvent in *vacuo* followed by column chromatography of the residue (silica gel, petroleum ether -EtOAc) furnished **N*b*B**, which was recrystallized from $CH_2Cl_2$/cyclohexane to afford orange needles (190 mg, 73%). $^1H$ NMR (600 MHz, $(CD_3)_2CO$, 25 °C, TMS; Figure S5): δ 2.40 (s, 3H), 2.48 (s, 3H), 2.52 (s, 3H), 2.53 (s, 3H), 2.76 (s, 3H), 6.32 (s, 1H), 7.32 (d, $J$ = 9.0 Hz, 1H), 7.53 (t, $J$ = 7.8 Hz, 1H), 7.68 (t, $J$ = 7.8 Hz, 1H), 8.00 (d, $J$ = 8.4 Hz, 1H), 8.35 (d, $J$ = 9.0 Hz, 1H), 8.48 (d, $J$ = 8.4 Hz, 1H), 9.59 (s, 1H); $^{13}C$ NMR (200 MHz, $CDCl_3$, 25 °C, TMS; Figure S6): δ 11.5, 13.7, 14.8, 16.8, 17.7, 108.3, 119.1, 120.7, 123.1, 125.3, 128.0, 129.3, 129.7, 129.8, 131.4, 132.4, 132.9, 133.8, 141.4, 142.8, 144.1, 145.5, 158.1, 162.8, 163.4 ppm; HRMS (ESI-TOF) m/z: $[M+K]^+$ Calcd. for $C_{25}H_{23}B_2F_4N_3O$: 518.1595; Found: 518.1604.

**Procedure for Dye Sensitized Photo-oxidation Study.** This experiment was carried out following a previous report with minor modifications. Breifly, the aerated ethanol solutions of DPBF (50 μM) and **BP**, **N*m*B** and **N*b*B** (5 μM each) in a 10 mm quartz cell was irradiated over a time period of 18 min, using a stablized tungsten lamp (intensity: 40 $W/m^2$) with a wavelength cut offfilter >495 nm. Under these conditions, only the BODIPY dyes were excited to generate $^1O_2$, which would primarily react with DPBF because the reaction rate of $^1O_2$ is significantly higher with DPBF than the BODIPY dyes and also, DPBF was taken at 50 times higher molar concentration in comparison to the BODIPY dyes. The absorbance of the solutions were measured at $\lambda_{abs}$ of DPBF (412 nm) at an interval of 1 min. The relative $^1O_2$ generation capacities of dyes **BP**, **N*m*B** and **N*b*B** in ethanol were determined by measuring the time-dependent reduction of the DPBF absorbance.

**Compound Stock Preparation.** Stock solutions (10 mM) of **N*m*B**, **N*b*B** and **BP** were prepared in DMSO, stored as aliquots at -20 °C and used with in 3 months of preparation. Final concentrations of all the tested compounds and other fluorescent probes were prepared in complete medium (DMEM+ 10% FBS). The final concentration of DMSO in all the samples was less than 0.1%.

**MTT Assay.** Cell viability of control and treated cells were assessed by the MTT reduction assay, as per the previous report[2] with minor modifications. Breifly, MIA PaCa-2 and PANC-1 cells were seeded (4.5 x $10^3$/well) in 96 well plate. After 24 h, the unsynchronised cells were treated with vehicle (0.1% DMSO) or different concentrations of the respective compounds for 30 min, followed by photo-irradiation for indicated time periods. After 48 h incubation (5% $CO_2$, 37°C). cells were washed with PBS, and 100 µl of 0.5 mg/ml- MTT solution was added and incubated for 3 h to form formazan crystals. Later formazan crystals were solubilized in DMSO, and colour intensity was recorded using a multiplate reader ($\lambda_{abs}$ = 570 nm; Polestar, BMG Labtech). Percentage cell viability was calculated and plotted against untreated control.

**Cellular Thiol Estimation Assay.** Estimation of thiol content was determined by Flow cytometer after staining cells with MBB (Monobromobimane), following a published protocol. Briefly, cells were seeded in 6 well plates (6.0 x $10^4$ cells/well) for overnight, treated with **N*b*B** (5 µM) and irradiated with visible light (30 min). After 3 h and 24 h of incubation, cells were stained with MBB (40 µM) for 30 min, washed twice with PBS, collected by trypsinization and again washed twice. Fluorscence of the cells were acquired using Partec CyFlow flow cytometer. At least 25,000 cells per sample were acquired and analyzed by FlowJo software.

**Western Blotting.** Western blotting analysis was done as per the standard procedure, as reported previously. Briefly, cells were seeded in culture dish (1.25 x $10^6$ cells/dish) for overnight, treated with **N*b*B** (0-15 µM) and irradiated with visible light (30 min). After 24 h of incubation, cells were harvested by scraping and washed twice with PBS. Cells were lysed with lysis buffer (20 mM HEPES, 0.5 mM EGTA, 2 mM EDTA, 1 mM DTT, 250 mM NaCl, 1% NP-40) supplemented with protease and phosphatase inhibitors cocktail. After estimating proteins with BCA Protein Assay Kit (Pierce™), 50 µg cell lysate protein was resolved into bands on 10% SDS- PAGE. Proteins in the gels were transferred to nitrocellulose membrane and the membrane was blocked using PBST (PBS - 0.1% Tween 20) supplemented with 2% non-fat milk for 2 h. Blots were incubated with primary antibodies in PBST containing 1% non-fat milk for overnight at 4 °C. Blots were washed thrice in PBST (10 min each) and incubated with secondary antibody conjugated to HRP for 3h. After washing thrice with PBST, blots were developed using Lumi-Light$^{plus}$ western blotting kit by Kodak Gel-doc.

Band image processing and band intensity analysis was done using Kodak Gel-doc, and the band intensity was estimated using ImageJ software, and then protein expression difference was calculated after normalization of the treated sample to the untreated sample.

### Supplementary Figures

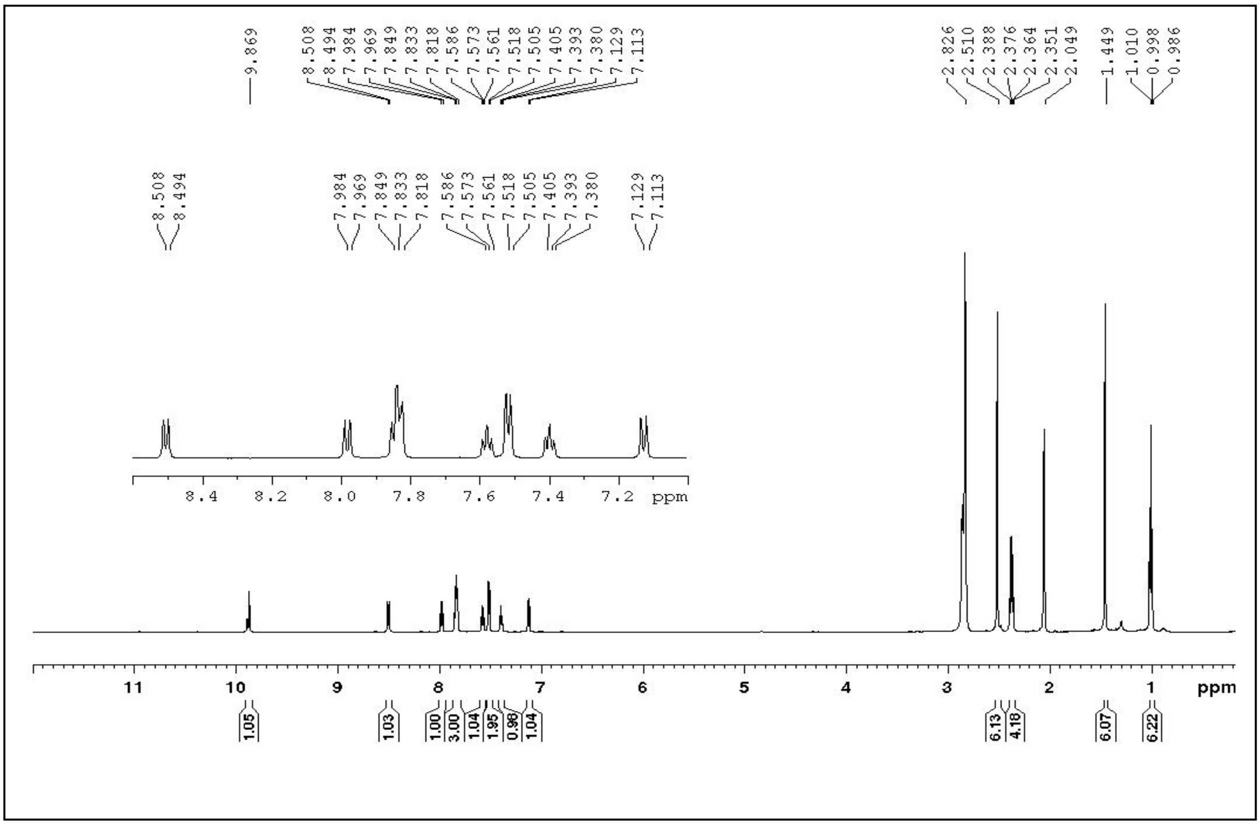


**Figure S1.** $^{1}$H NMR spectrum of dye **2** in $(CD_3)_2CO$ (600MHz).

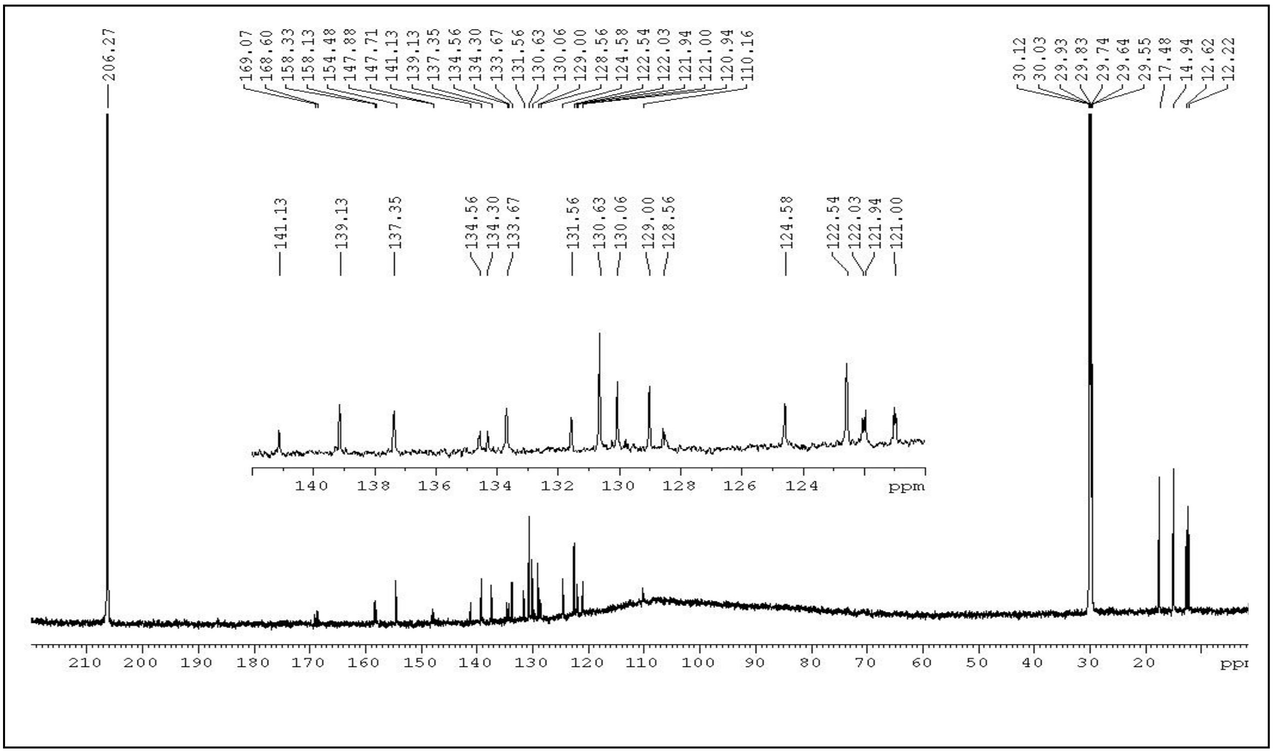


**Figure S2.** $^{13}$C NMR spectrum of dye **2** in $(CD_3)_2CO$ (800MHz)

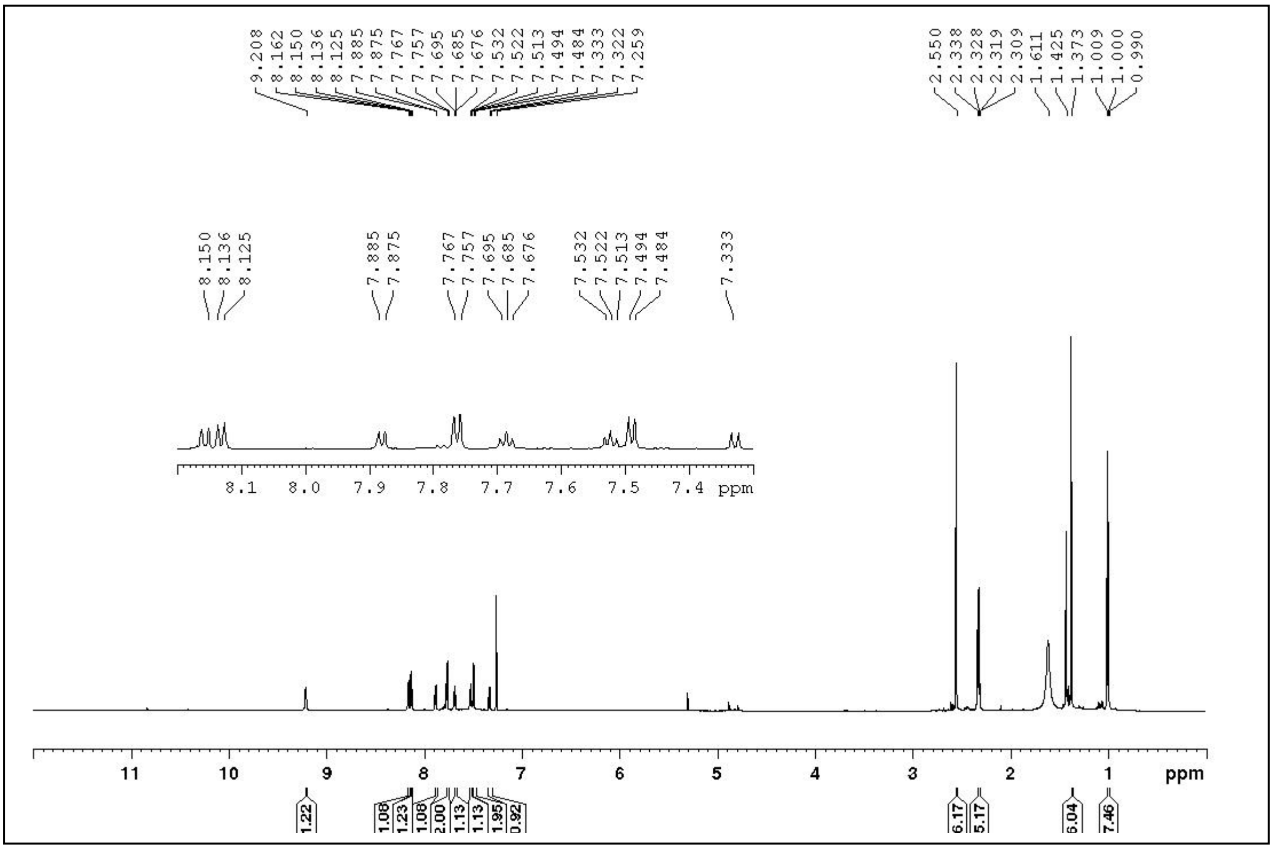


**Figure S3.** $^{1}$H NMR spectrum of **N*m*B** in $CDCl_3$ (800 MHz).

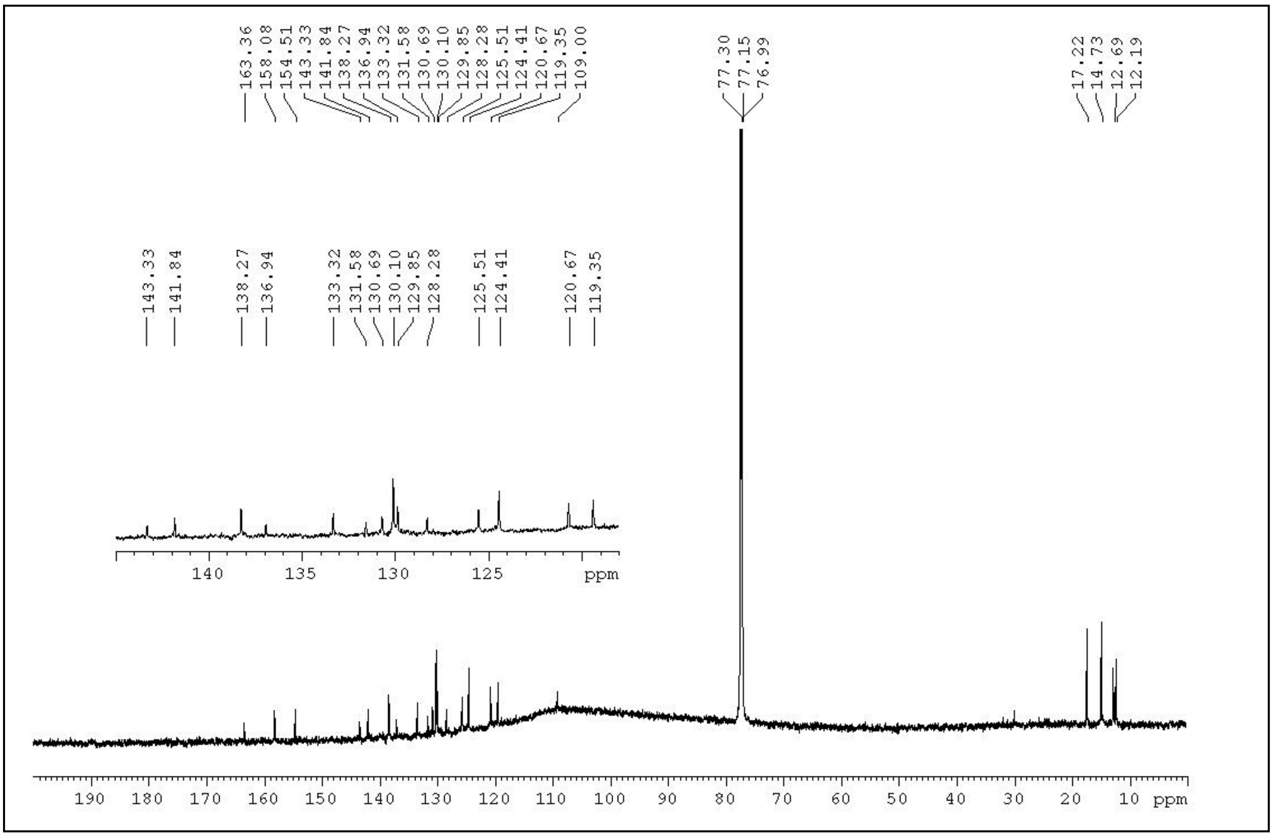


**Figure S4.** $^{13}$C NMR spectrum of **N*m*B** in $CDCl_3$ (800 MHz).

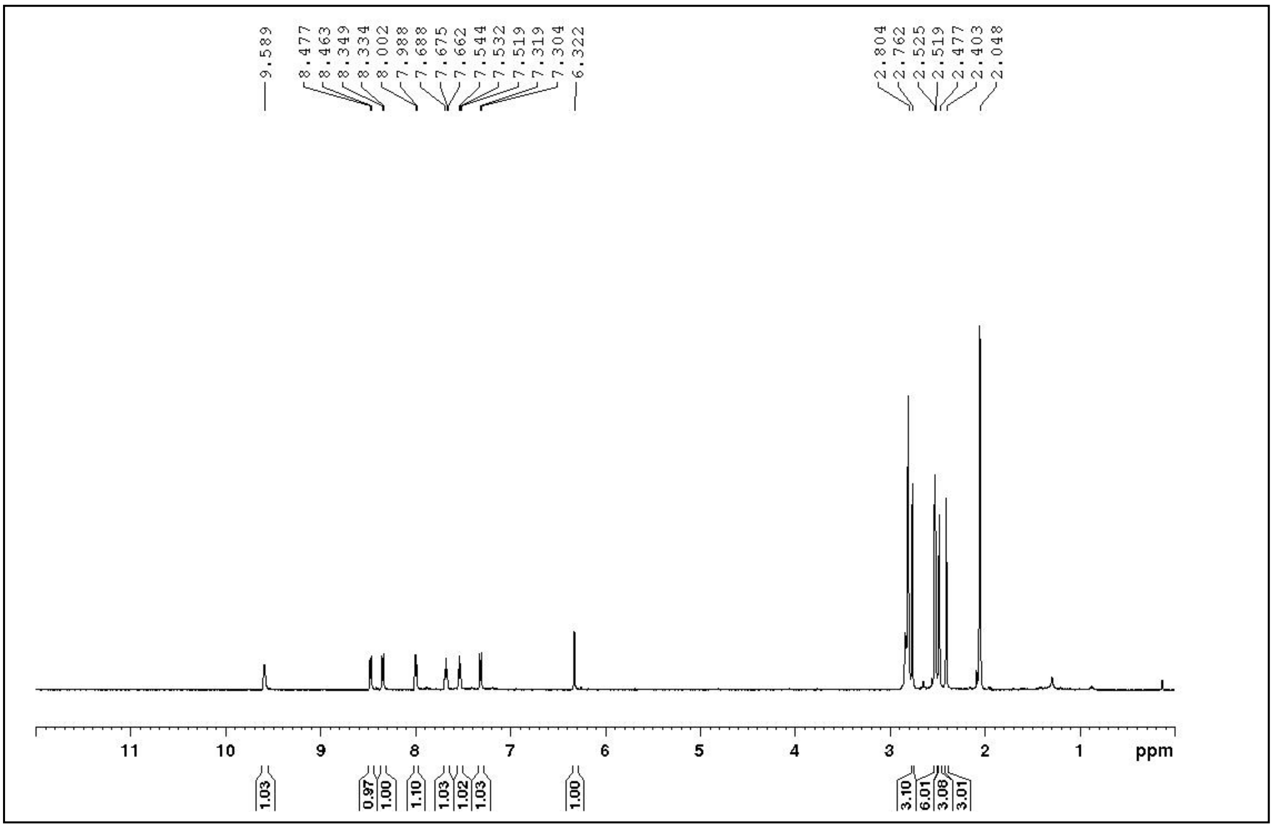


**Figure S5.** 1H NMR spectrum of **NbB** in (CD3)2CO (600 MHz).

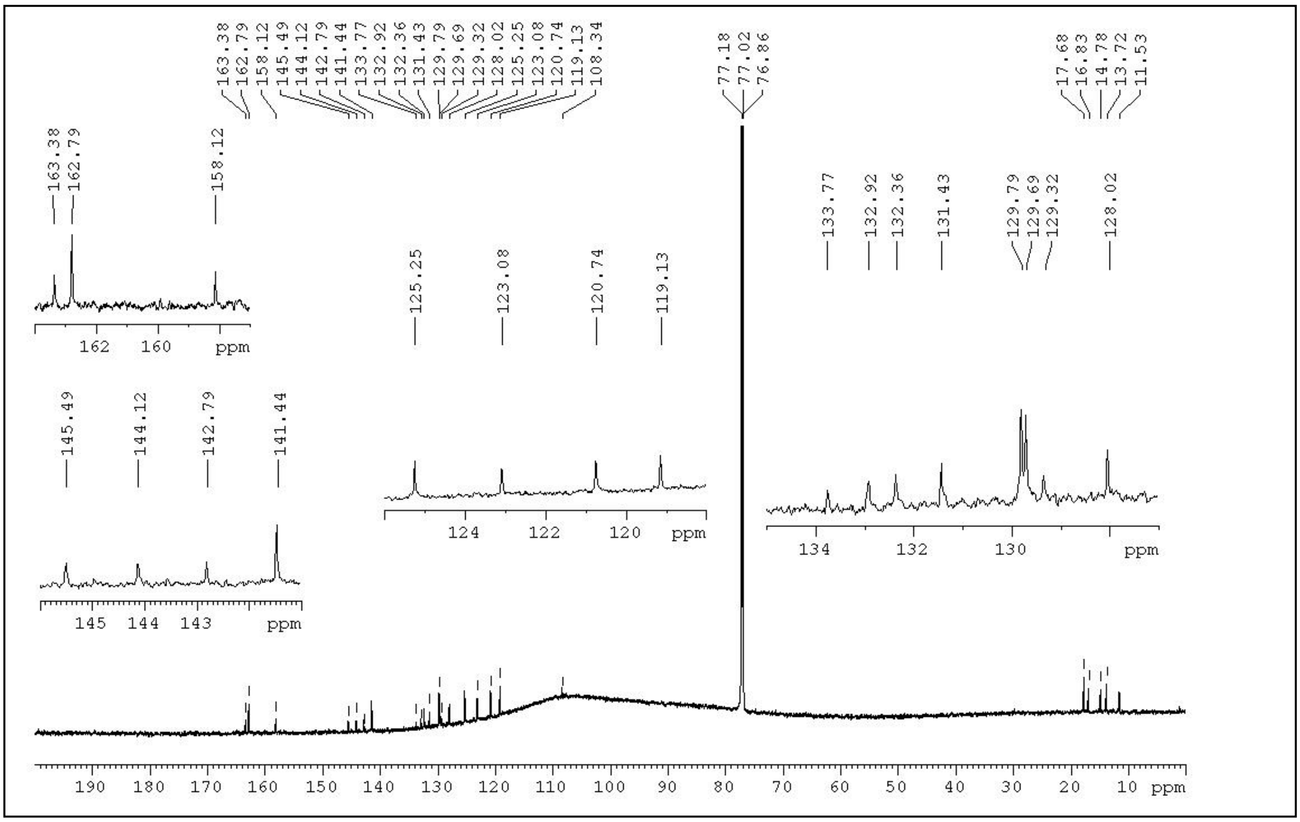


**Figure S6.** $^{13}$C NMR spectrum of **N*b*B** in $CDCl_3$ (800 MHz).

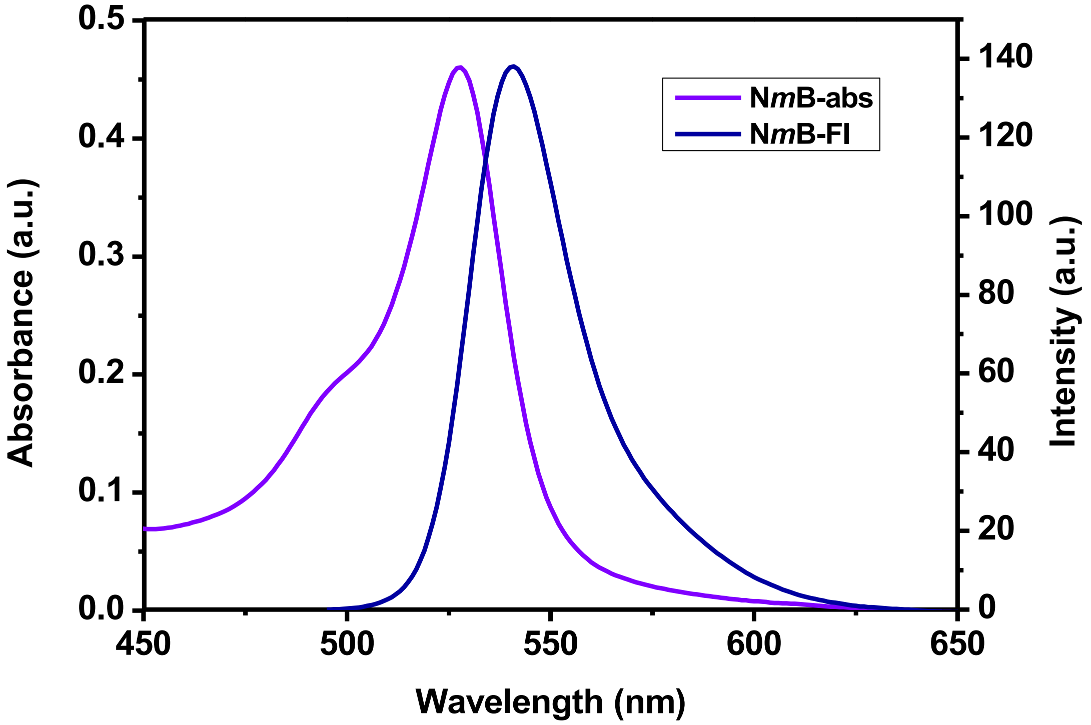


**Figure S7.** Absorption and emission spectra of ***NmB*** in $CH_2Cl_2$.

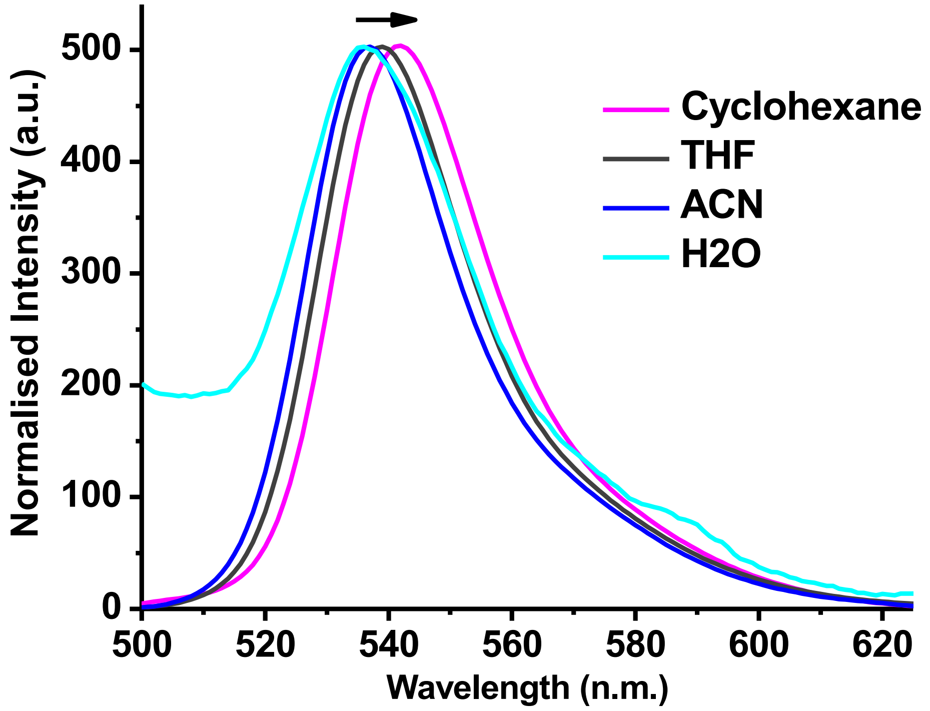


**Figure S8.** Fluorescence spectra of **N*m*B** were acquired in different solvents.

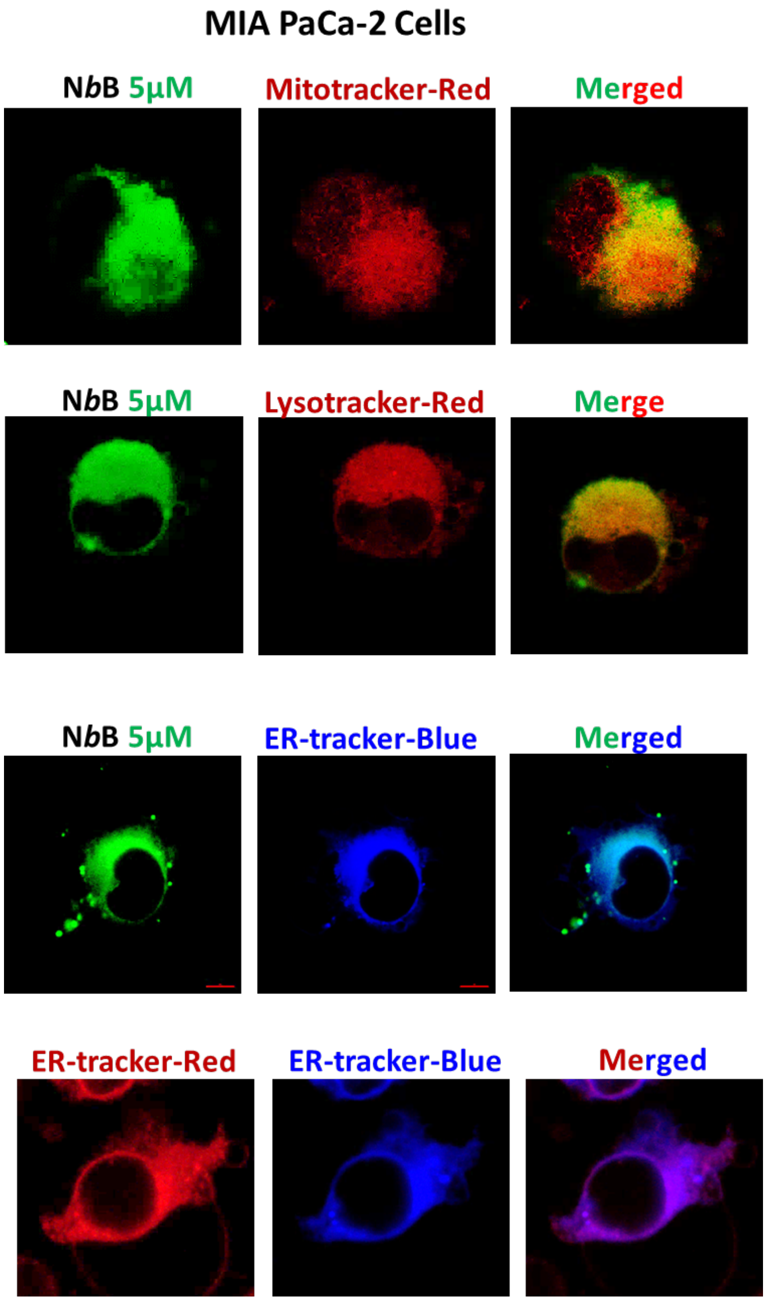


**Figure S9. Colocalization of NbB with Mito Tracker-Red, Lyso Tracker-Red and ER Tracker-Blue.** MIA PaCa-2 cells were coincubated with **NbB** (5 µM) and Mito Tracker-Red, Lyso Tracker-Red or ER Tracker-Blue cells for 30 min. In another set, MIA PaCa-2 cells were coincubated with ER Tracker-Blue and ER Tracker-Red for 30 min. Fluoroscnce images were acquired in multi-channel mode in confocal microscope. (Channel green: $\lambda_{ex}$ = 488 nm; Channel red: $\lambda_{ex}$ = 594 nm; Channel Blue: $\lambda_{ex}$ = 355 nm).

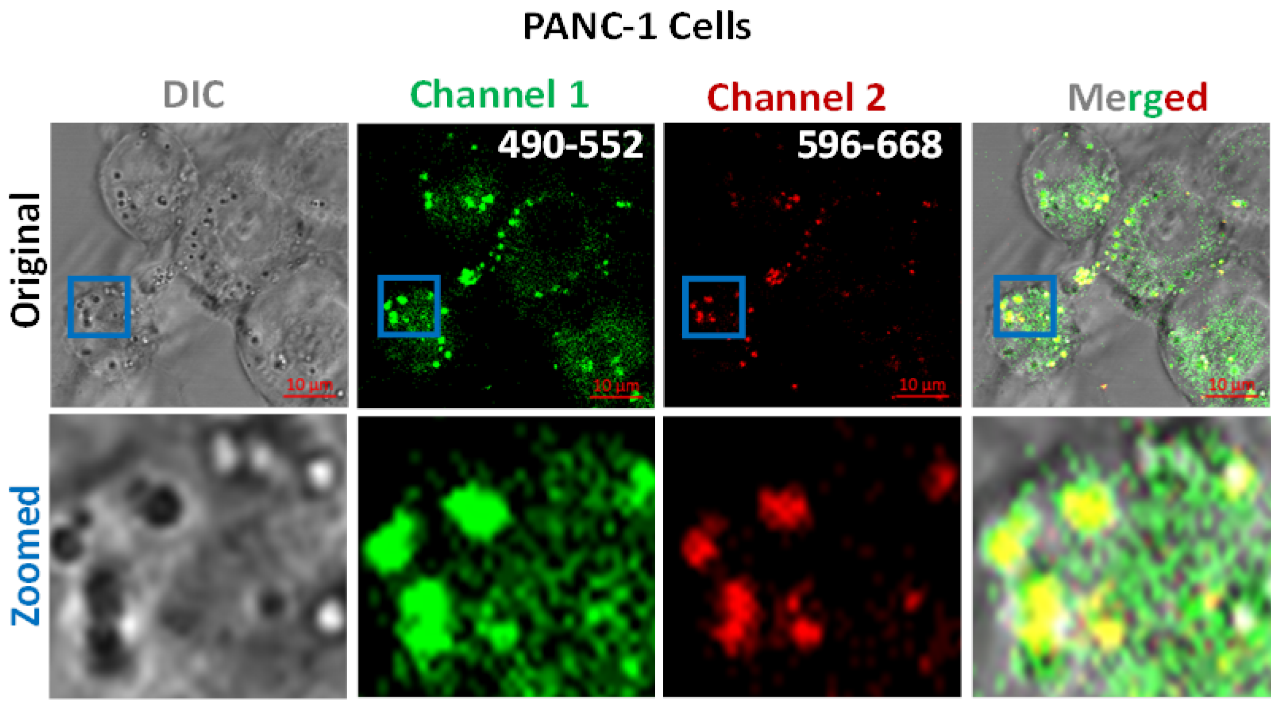


**Figure S10.** PANC-1 cells were treated with **N*b*B** (5 µM each) for 30 min and DIC and fluoroscnce images were acquired in multi channel mode confocal microscope. Zoomed images show localization of **N*b*B** in ER (green only) and LD (yellow colour due to colocalization of geen in channel 1, red in channel 2 and black in DIC channel for globular lipid droplets). (Channel 1: $\lambda_{ex}$ = 488 nm; Channel 2: $\lambda_{ex}$ = 594 nm). Emission range was shown in "nm" in Channel 1 and 2.

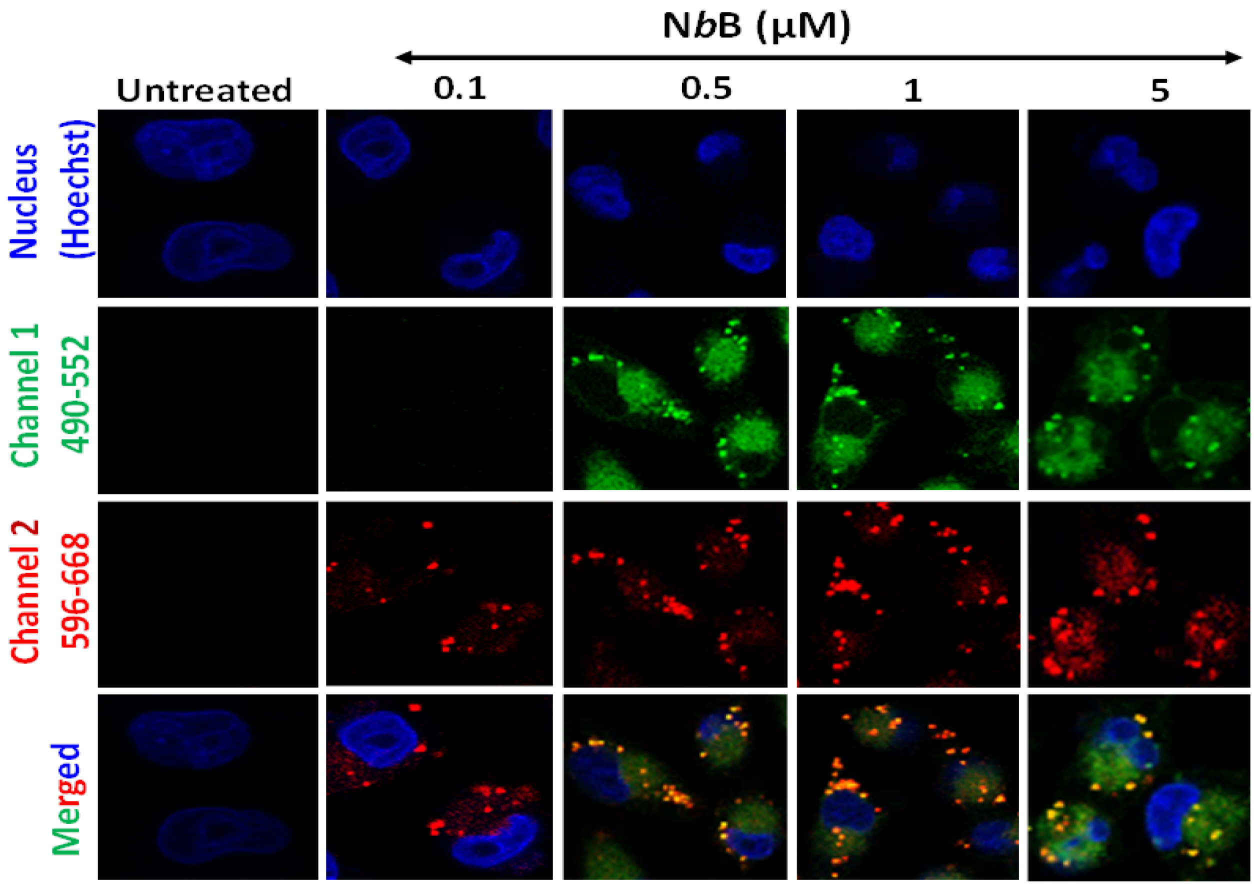


**Figure S11.** MIA PaCa-2 cells were treated with indicated concentrations of **N*b*B** for 30 min and fluoroscnce images were acquired in multi channel mode in confocal microscope. (Channel 1: $\lambda_{ex}$ = 488 nm; Channel 2: $\lambda_{ex}$ = 594 nm). Emission range was shown in "nm" in Channel 1 and 2.

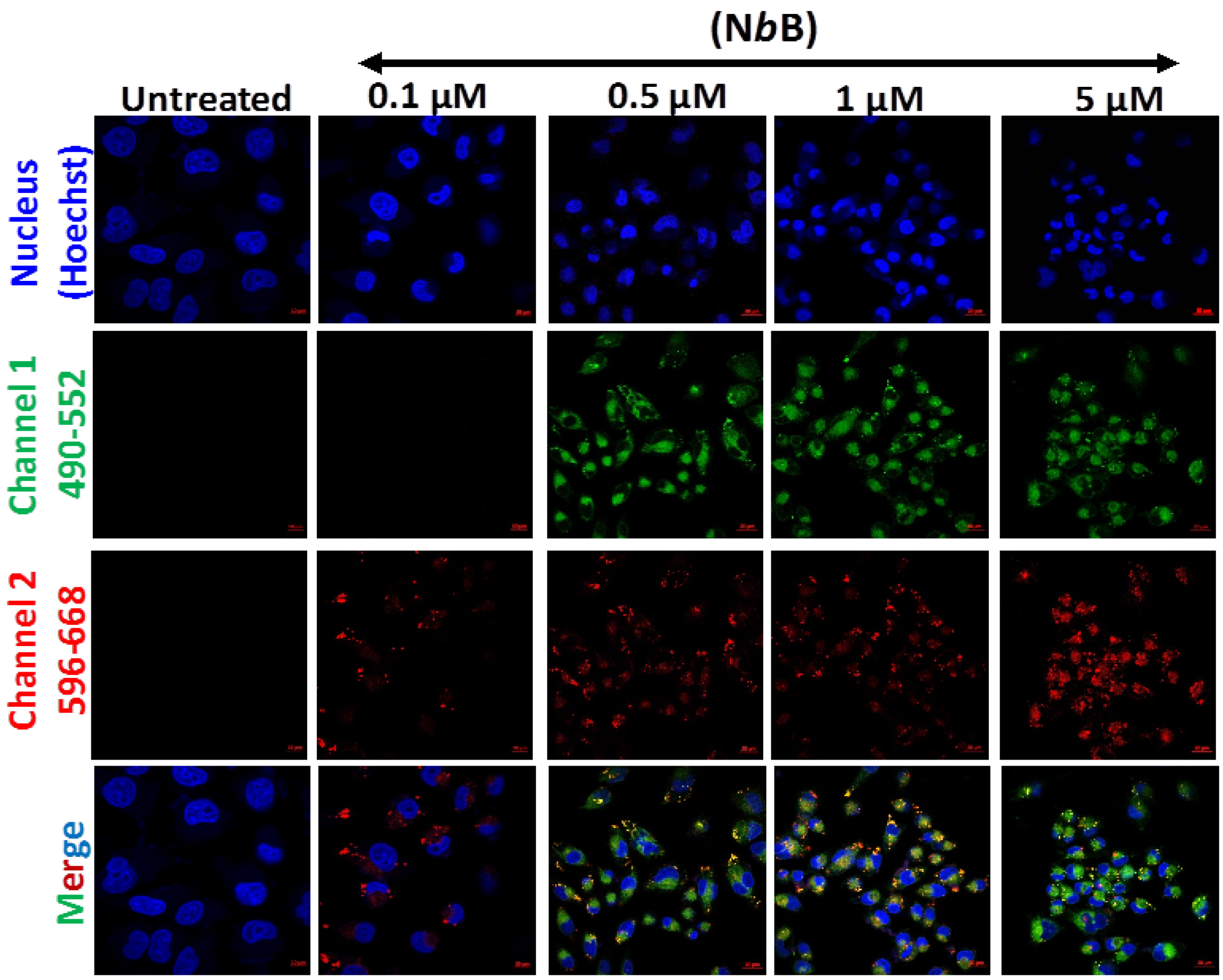


**Figure S12.** PANC-1 cells were treated with indicated concentrations of **N*b*B** for 30 min and fluoroscence images were acquired for multiple cells in multi channel mode in confocal microscope. (Channel 1: $\lambda_{ex}$ = 488 nm; Channel 2: $\lambda_{ex}$ = 594 nm). This experiment showed uniform labelling of the **N*b*B** in all the cells.

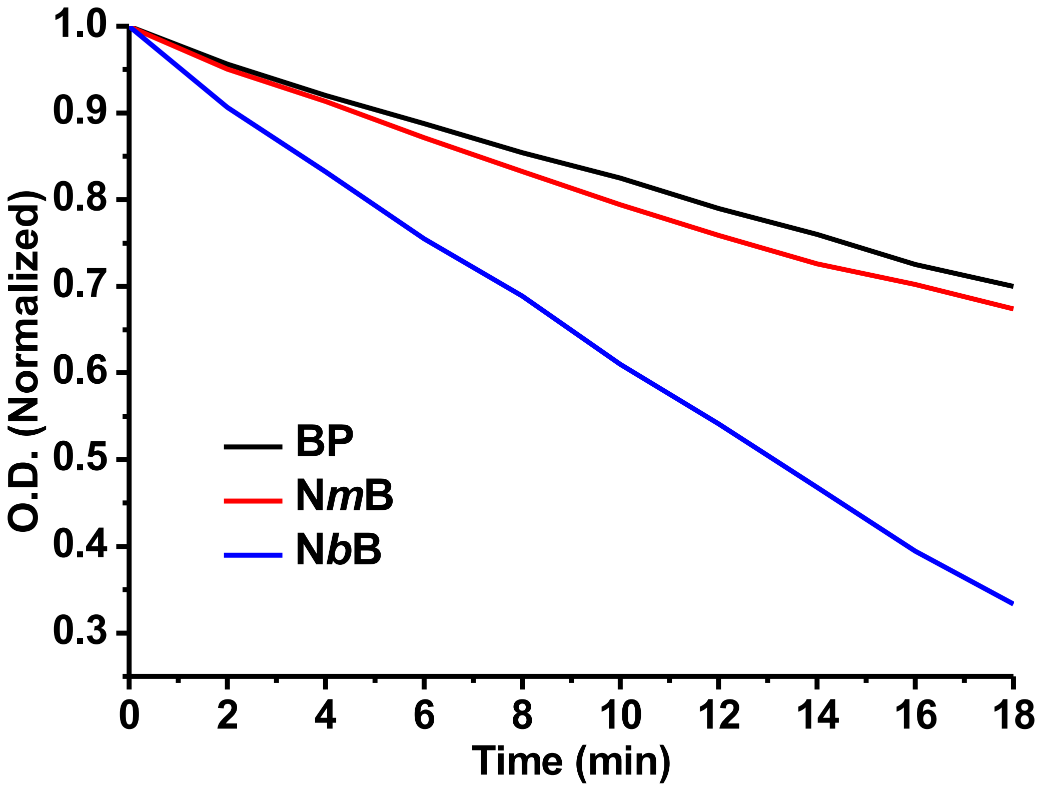


**Figure S13.** Time-dependent depletion of DPBF (50 μM) absorbance at $\lambda_{max}$ (412 nm) under dye-sensitized photooxidation in the presence of **BP**, **N*m*B** and **N*b*B** (5 μM) in ethanol.

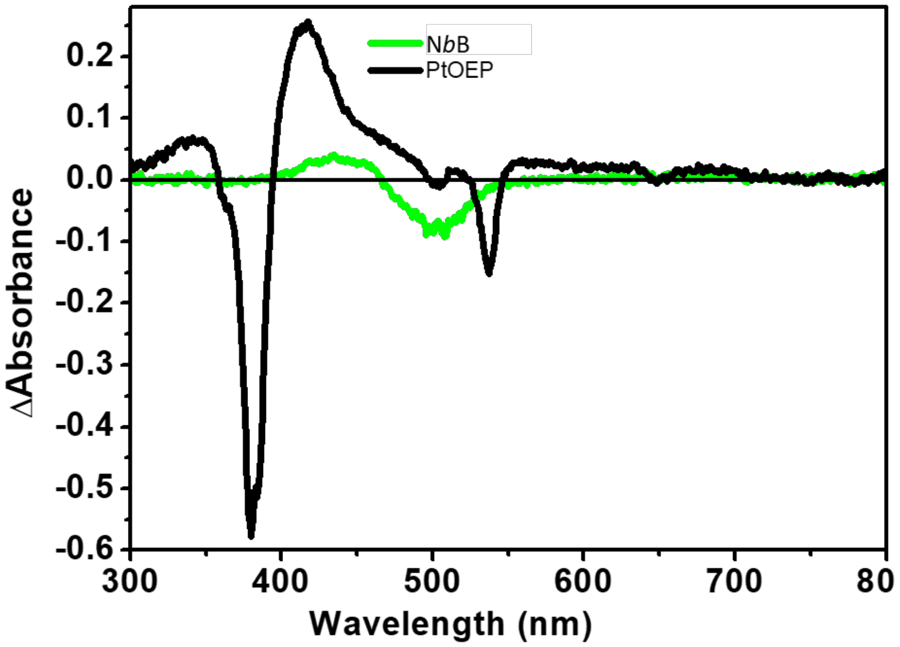


**Figure S14.** Comparative measurement of triplet spectrum of **N*b*B** and Platinum octaethylporphyrin (triplet yield 1) for triplet yield determination. Experiments were performed at identical experimental condition (same sample OD and excitation energy at excitation wavelength of 532 nm).

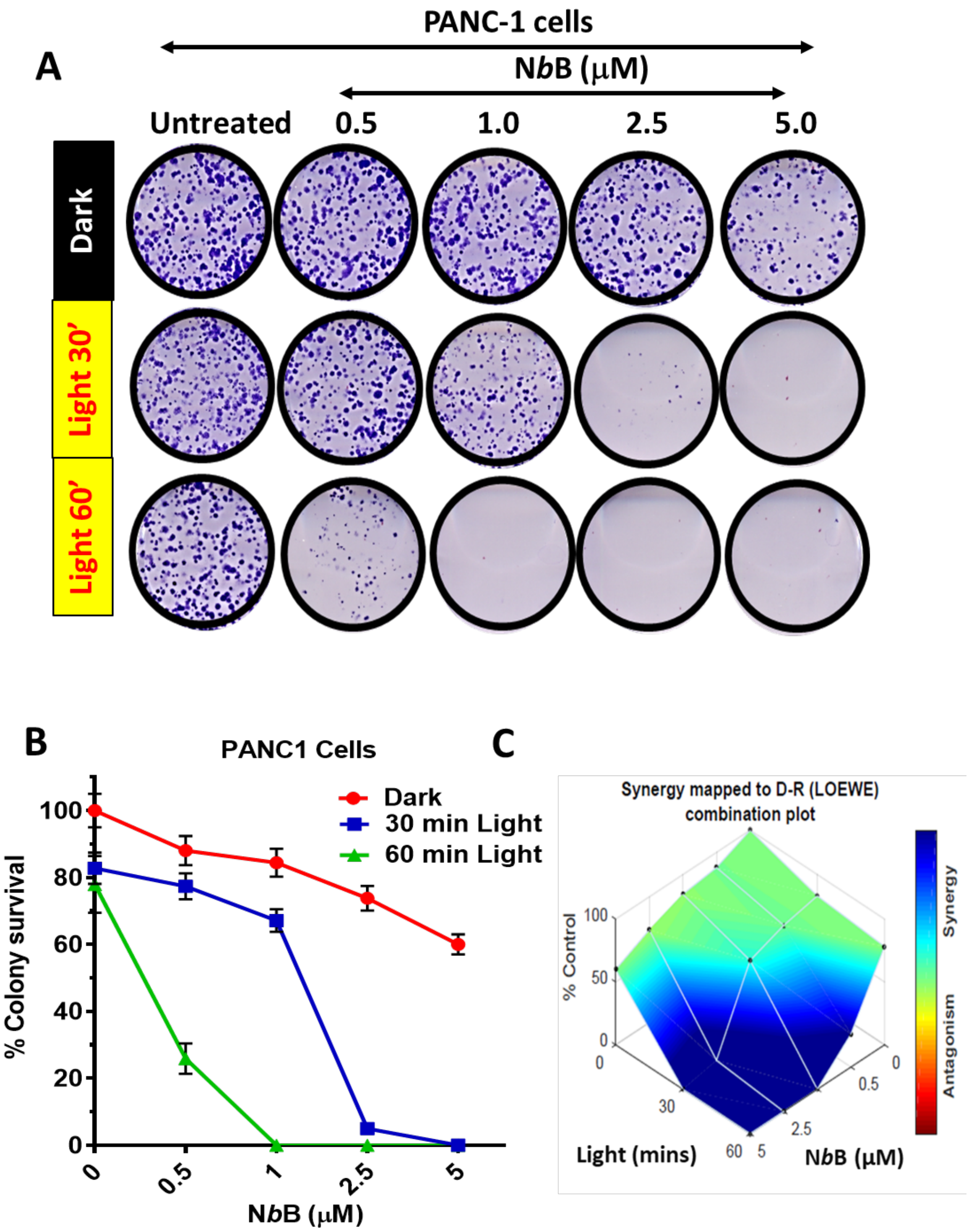


**Figure S15.** (A) PANC-1 cells were treated with indicated concentrations of **N*b*B** for 30 min and photoirradiated for 0 (dark)/30/60 min. Cells after allowing to grow colonies for 9-10 days, colonies were stained with crystal violet. (B) Quantification of % colony survival from the above experiment. (C) Synergistic interaction of **N*b*B** and light in PDT effects in the above experiment.

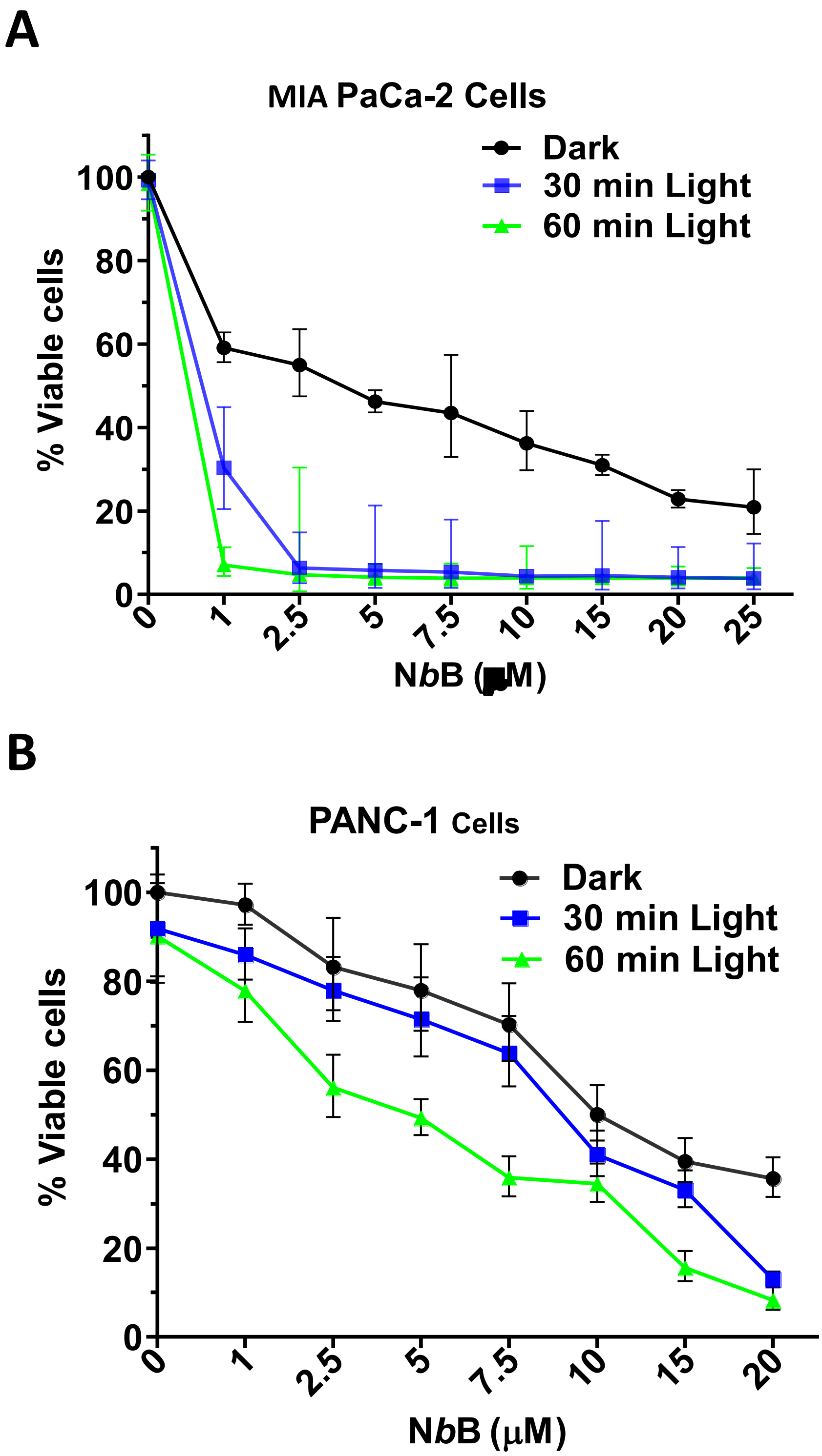


**Figure S16.** (A, B) MIA-PaCa-2 and PANC-1 cells were treated with indicated concentrations of **N*b*B** for 30 min and photoirradiated for 0 (dark)/30/60 min. Cell viability was assessed by MTT assay after 48 h.

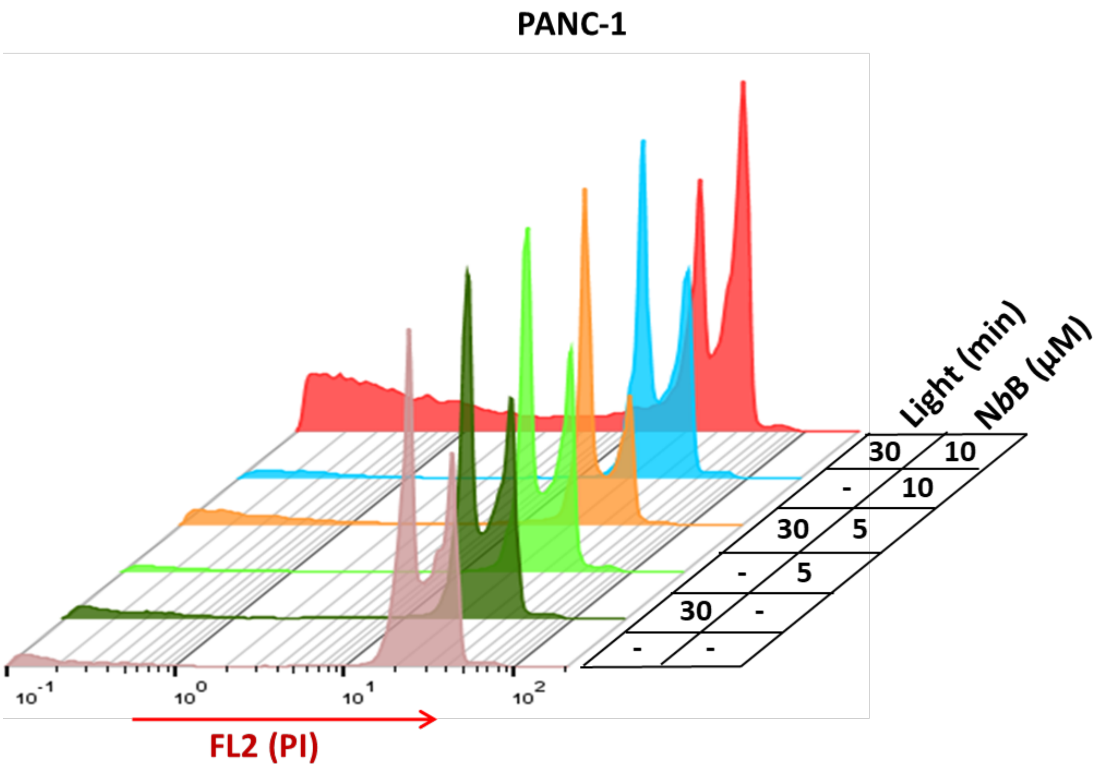


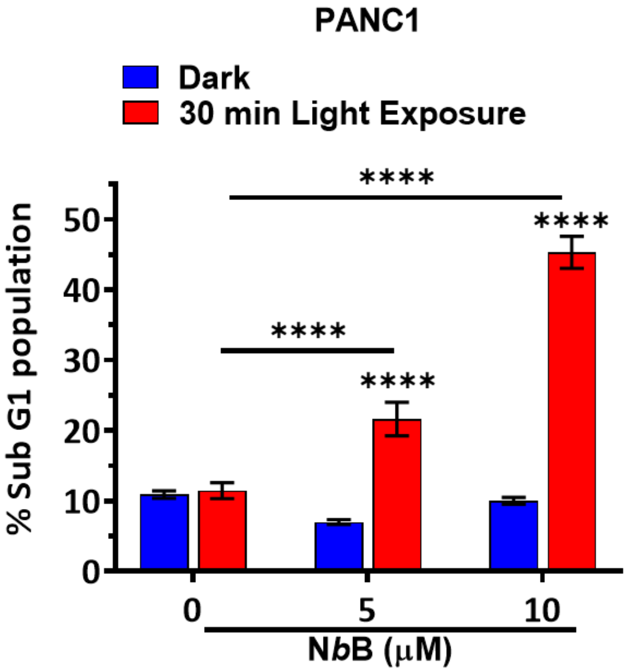


**Figure S17.** PANC-1 cells were treated with indicated concentrations of **N*b*B** for 30 min and photoirradiated for 0 (dark)/30 min. Sub-G1 analysis was carried out after 24 h.

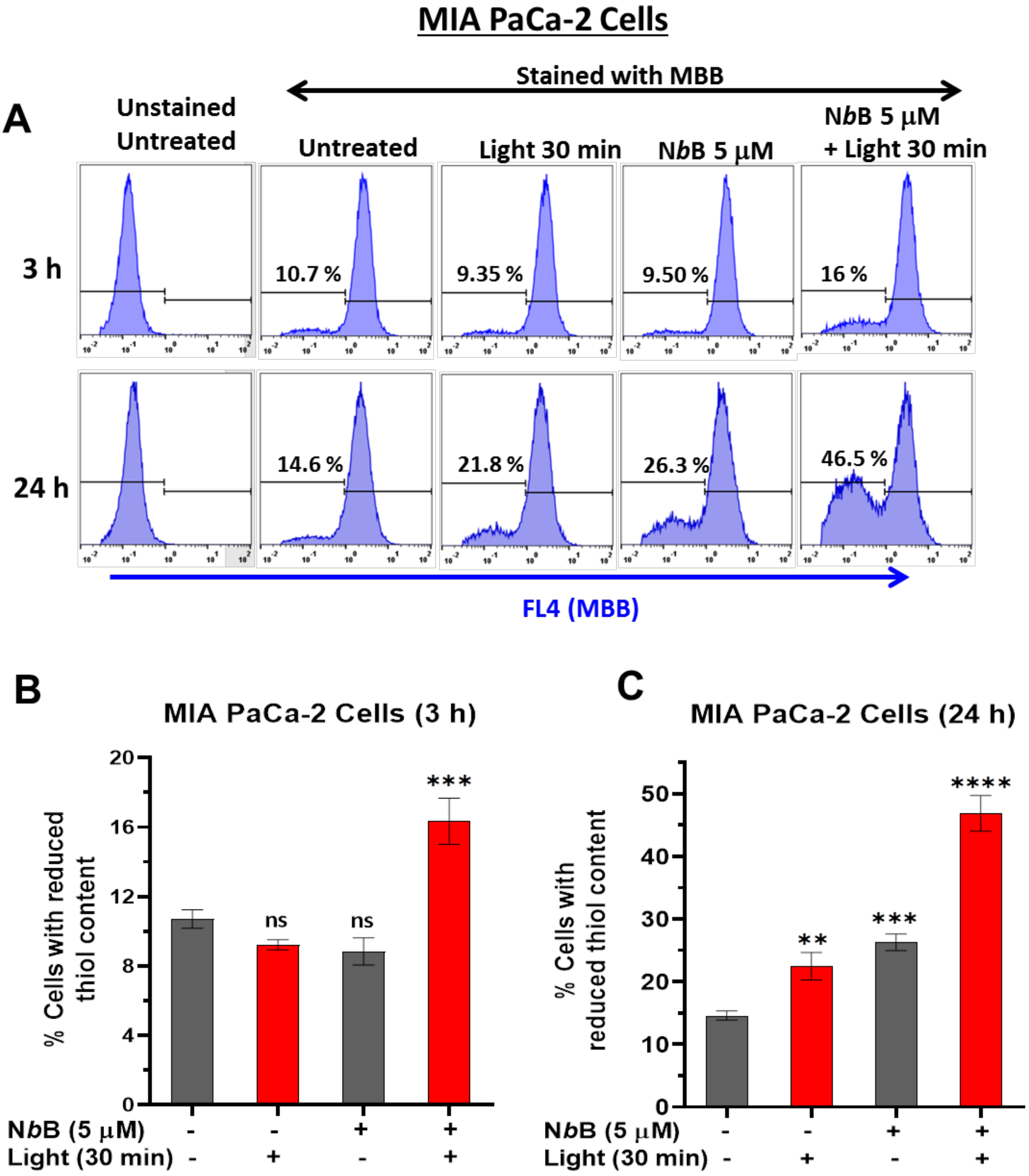


**Figure S18.** (A) MIA-PaCa-2 cells were treated with indicated concentrations of **N*b*B** for 30 min, photoirradiated for 0 (dark)/30 min and incubated for 3 /24 h. Total cellular thiol content was measured using MBB dye by flow cytometry. (B, C) Quantification of cells with reduce thiol contents, at 3 h or 24 h recovery, in athe above experiments. **$p$<0.01, ***$p$<*.001*, ****$p$<0.0001 and ns: not significantly different w.r.t untreated control.

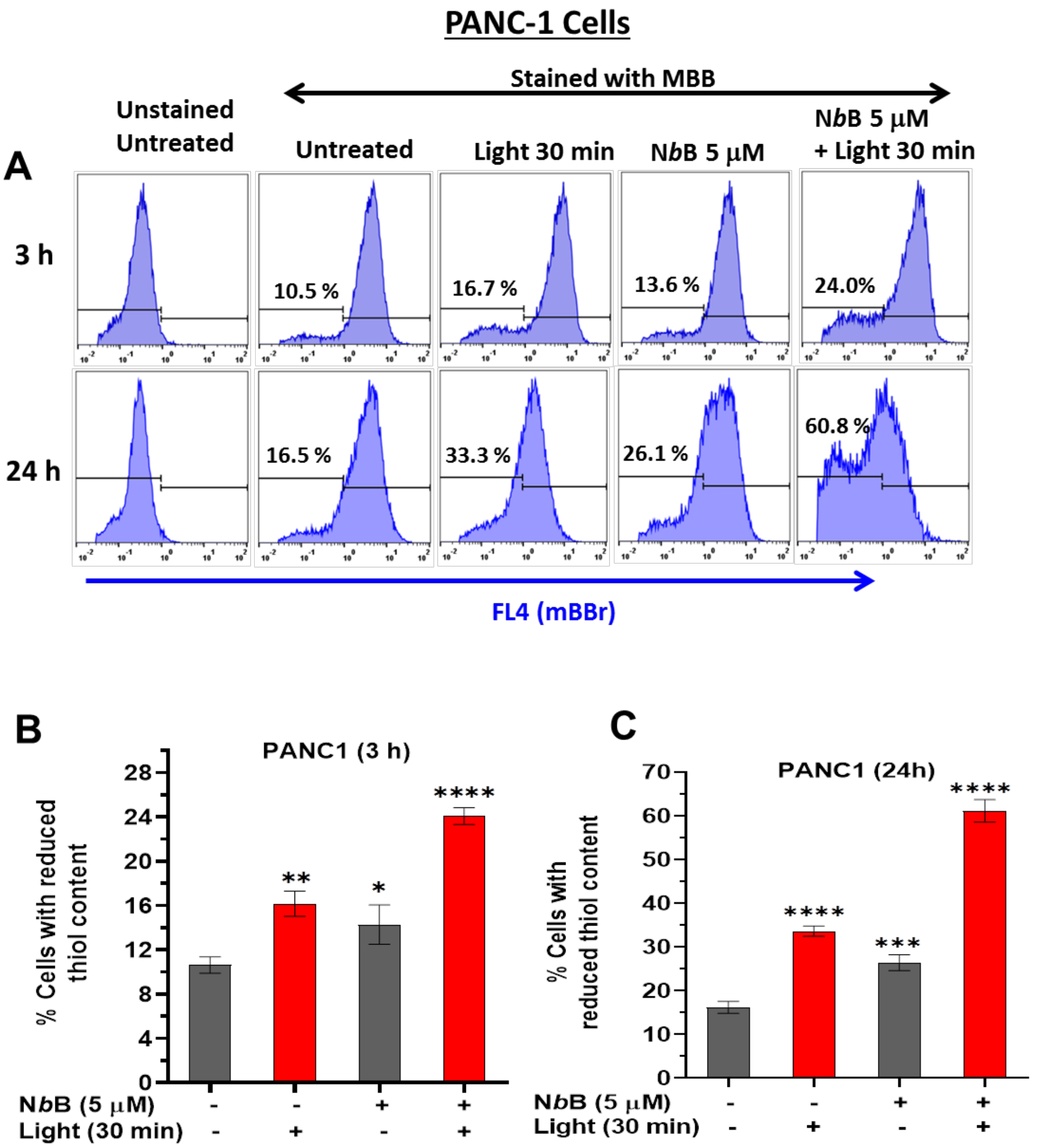


**Figure S19.** (A) PANC-1 cells were treated with indicated concentrations of **N*b*B** for 30 min, photoirradiated for 0 (dark)/30 min and incubated for 3 /24 h. Total cellular thiol content was measured using MBB dye by flow cytometry. (B, C) Quantification of cells with reduce thiol contents, at 3 h or 24 h recovery, in the above experiments. **p*<0.05, ***p*<0.01, ****p*<0.001, *****p*<0.0001 w.r.t untreated control.

### Supplementary Tables

**Table S1.** Photophysical parameters of the dyes **N*m*B**, **BP** and **N*b*B** in DCM.

| Dye | $\lambda_{abs}$ [nm] | $\varepsilon_{max}$ [$M^{-1}cm^{-1}$] | $\lambda_{em}$ [nm] | $\nu$ [$cm^{-1}$] | $\Phi_{fl}$ | τ (ns) |
|---|---|---|---|---|---|---|
| **N*m*B** | 528.0 | 18479 | 543.0 | 523 | 0.04[a] | 5.4 |
| **BP**[3] | 493.0 | 79000 | 504.0 | 443 | 0.99 | 3.2 |
| **N*b*B** | 502.0 | 24939 | 541.4 | 1436 | 0.09[a] | 0.6 |

[a] Determined using $\Phi_{fl}$ = 0.88 for Rhodamine 6G in ethanol as the reference. [1]

**Table S2.** Photophysical parameters of the **N*b*B** in different solvents.

| Solvent | $\lambda_{abs}$ [nm] | $\varepsilon_{max}$ [$M^{-1}cm^{-1}$] | $\lambda_{em}$ [nm] | $\nu$ [$cm^{-1}$] |
|---|---|---|---|---|
| **Cyclohexane** | 506 | 30367 | 544 | 1381 |
| **THF** | 502 | 30297 | 539 | 1367 |
| **ACN** | 497 | 23215 | 525 | 1073 |
| **$H_2O$** | 501 | 1957 | 514 | 505 |

**Table S3.** Photophysical parameters of the **N*m*B** in different solvents.

| Solvent | $\lambda_{abs}$ [nm] | $\varepsilon_{max}$ [$M^{-1}cm^{-1}$] | $\lambda_{em}$ [nm] | $\nu$ [$cm^{-1}$] |
|---|---|---|---|---|
| **DCM** | 528 | 15780 | 541 | 455 |
| **Cyclohexane** | 528 | 17580 | 542 | 489 |
| **THF** | 525 | 11670 | 539 | 495 |
| **ACN** | 523 | 16890 | 537 | 498 |
| **$H_2O$** | 530 | 15400 | 536 | 211 |
